# Cell-Type Resolved Protein Atlas of Brain Lysosomes Identifies SLC45A1-Associated Disease as a Lysosomal Disorder

**DOI:** 10.1101/2024.10.14.618295

**Authors:** Ali Ghoochani, Julia C. Heiby, Eshaan S. Rawat, Uche N. Medoh, Domenico Di Fraia, Wentao Dong, Marc Gastou, Kwamina Nyame, Nouf N. Laqtom, Natalia Gomez-Ospina, Alessandro Ori, Monther Abu-Remaileh

## Abstract

Mutations in lysosomal genes cause neurodegeneration and neurological lysosomal storage disorders (LSDs). Despite their essential role in brain homeostasis, the cell-type-specific composition and function of lysosomes remain poorly understood. Here, we report a quantitative protein atlas of the lysosome from mouse neurons, astrocytes, oligodendrocytes, and microglia. We identify dozens of novel lysosomal proteins and reveal the diversity of the lysosomal composition across brain cell types. Notably, we discovered SLC45A1, mutations in which cause a monogenic neurological disease, as a neuron-specific lysosomal protein. Loss of SLC45A1 causes lysosomal dysfunction in vitro and in vivo. Mechanistically, SLC45A1 plays a dual role in lysosomal sugar transport and stabilization of V1 subunits of the V-ATPase. SLC45A1 deficiency depletes the V1 subunits, elevates lysosomal pH, and disrupts iron homeostasis causing mitochondrial dysfunction. Altogether, our work redefines SLC45A1-associated disease as a LSD and establishes a comprehensive map to study lysosome biology at cell-type resolution in the brain and its implications for neurodegeneration.

## Introduction

Lysosomes are membrane-bound organelles responsible for degrading macromolecules and clearing damaged organelles to maintain cellular homeostasis^1,2^. Lysosomal function is mediated by its protein constituents, including lumenal hydrolases that degrade diverse cargo delivered from both intra- and extracellular sources, and integral and membrane-associated proteins that regulate catabolite export, lysosomal integrity, trafficking, fusion processes, and pH^3^.

Beyond degradation, lysosomes are key regulators of nutrient and energy sensing pathways, including mTORC1 and AMPK^1,2^, and are integral to the cell’s metabolic state^4,5^. Lysosomes serve as reservoirs for nutrients, such as amino acids, cholesterol, and metals like calcium and iron^6–9^. Consequently, they are involved in functions such as plasma membrane repair, stress resistance, cell growth, and programmed cell death^1,2^.

Given their central roles, mutations in lysosomal genes cause lysosomal storage disorders (LSDs) and contribute to a wide range of diseases, in particular neurodegenerative disorders such as Alzheimer’s disease, Parkinson’s disease, and frontotemporal dementia^5,10,11^. LSDs are marked by the accumulation of undigested macromolecules in lysosomes, and many of which manifest with severe neurological symptoms^12^.

While lysosomes function across tissues, the cell-type-specific composition and roles of lysosomes in the brain remain poorly understood. It is unclear whether lysosomal protein abundance varies between brain cell types and how this variation contributes to disease. Furthermore, it is unknown whether distinct lysosomal proteins drive specialized functions in different brain cells.

To address these gaps, we generated the first lysosomal protein atlas across the major brain cell types, neurons, astrocytes, oligodendrocytes, and microglia, using our LysoTag mouse model coupled with cell-type-specific Cre-recombinase expression^13^. This quantitative atlas reveals both shared and unique lysosomal proteins across brain cells. Notably, we identified SLC45A1, mutations in which cause a monogenic intellectual developmental disorder with neuropsychiatric features^14–16^, as previously unknown neuron-specific lysosomal protein involved in sugar transport. Mechanistic studies showed that SLC45A1 stabilizes the V-ATPase complex on the lysosomal membrane, and its loss leads to elevated lysosomal pH, disrupted iron homeostasis, and mitochondrial dysfunction—hallmarks of lysosomal storage disorders. These findings redefine SLC45A1-associated disease as a novel LSD. Our findings offer unprecedented insights into the cell-type-specific proteomic landscape of the lysosome in the brain, paving the way for novel discoveries in lysosomal biology and its role in neurological diseases.

## Results

### Cell-type-specific lysosome isolation and proteomic analysis using the LysoTag mouse

To isolate intact lysosomes from the four major brain cell types—neurons, astrocytes, oligodendrocytes, and microglia, we utilized our transgenic mouse model encoding the TMEM192–3xHA fusion protein (LysoTag) integrated into the *Rosa26* locus downstream of a lox-stop-lox (LSL) cassette^13^ (Figure 1A). By crossing the LysoTag mice with those expressing Cre recombinase under cell-type-specific promoters: *Syn1* for neurons^17,18^, *Gfap* for astrocytes^19,20^, *Olig2* for oligodendrocytes^21^, and *Cx3cr1* for microglia^22,23^, we generated mice that express the LysoTag in each brain cell type (Figure 1B). We confirmed that the LysoTag correctly localized to lysosomes in each cell type (Figures 1B, S1A). We then established and validated an efficient LysoIP protocol for the isolation of cell-type specific lysosomes (Figure S1B)^7,13^. Building on these results, we determined the proteomic landscape of purified lysosomes and the corresponding whole brain input lysates using label free data independent acquisition (DIA)-based mass spectrometry. As expected, we observed significant enrichment of known lysosomal proteins in the immunoprecipitates (IPs) from all the Cre lines compared to those from mockIP, derived from a mouse that does not express the LysoTag (Figures 1C, S1C, Tables S1A, S1D). Altogether, these results demonstrate that the LysoIP using a cell-type specific LysoTag system enables the isolation of lysosomes from different brain cell types without the need for prior cell sorting. This approach thus provides a robust method for studying cell-type-specific lysosomal composition.

**Figure 1.**
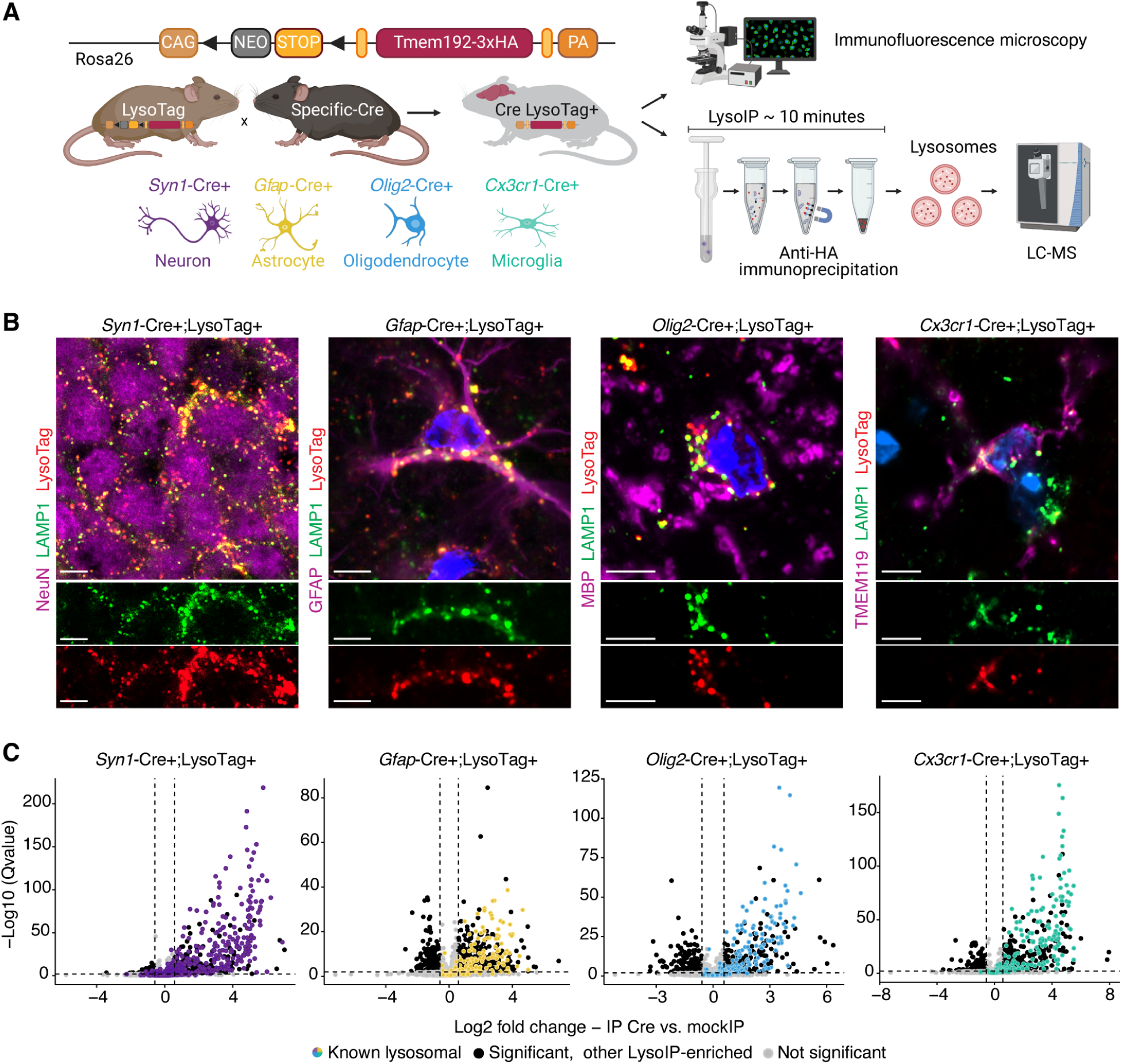
Cell-type-specific LysoIP from the brain. (A) Schematic representation of the experimental pipeline for isolating cell-type-specific lysosomes from mouse brains and further analysis by immunofluorescence microscopy of mass spectrometry (LC-MS). (B) Immunofluorescence staining to assess the colocalization of LysoTag (TMEM192-3xHA) with LAMP1, a lysosomal marker. NeuN was used as a marker for neurons, GFAP for astrocytes, MBP for oligodendrocytes, and TMEM119 for microglia. Scale bars are 5 µm. (C) Volcano plot representation of the log2 fold change (FC) in the abundance of proteins in IP samples from different Cre lines against mockIP controls (*n* = 3-6). Dashed lines indicate the threshold used to select differentially abundant proteins (absolute log2 FC > 0.58 and Q-value < 0.05). Colored dots highlight known lysosomal proteins, while black dots show other significantly LysoIP-enriched proteins, Table S1D. Known lysosomal proteins defined in Table S1A.

**Figure S1.**
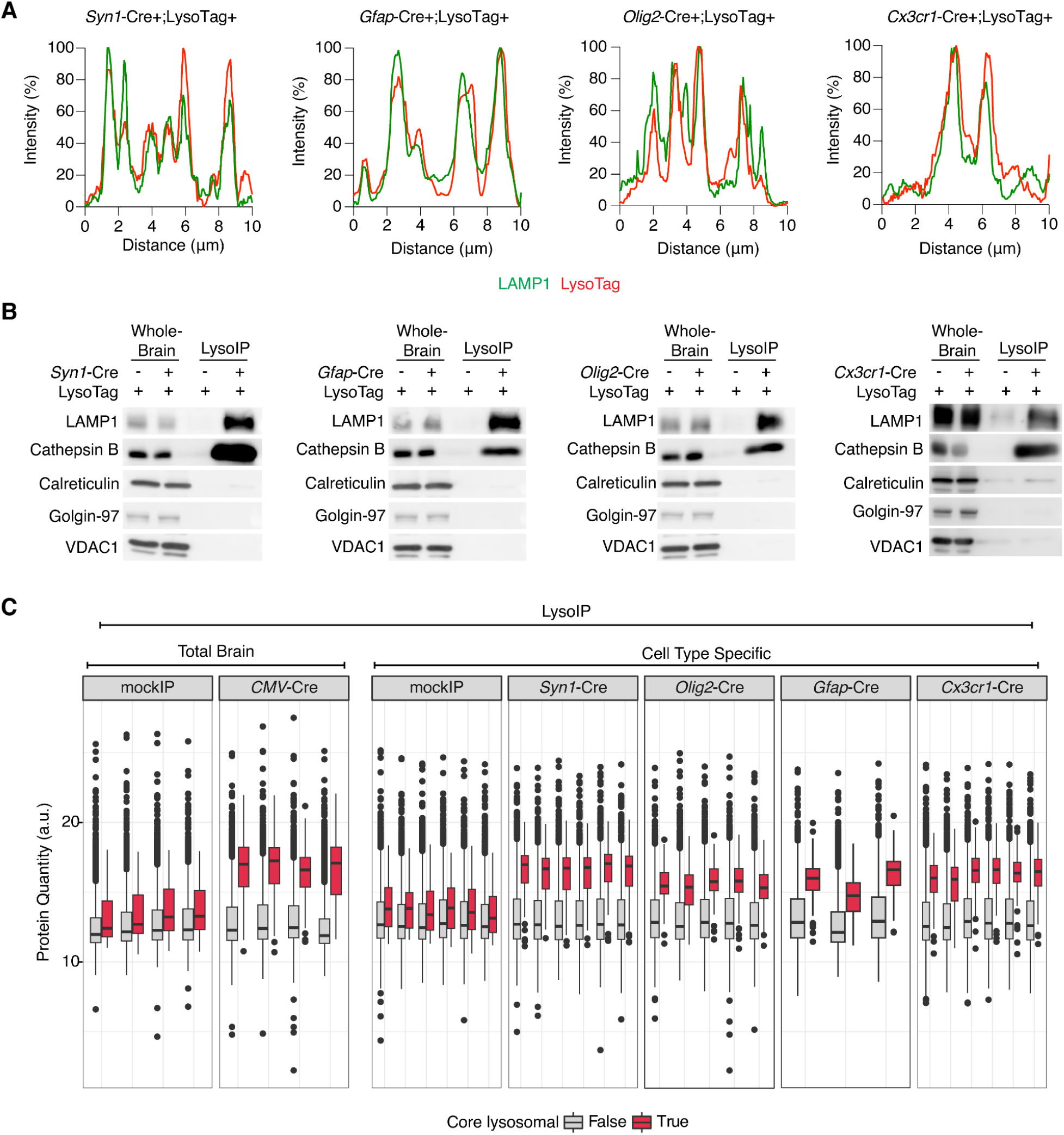
Validation and characterization of the cell-type LysoIP. (A) Pixel intensity plots for LAMP1 (Green) and LysoTag (Red) expression in neurons (*Syn1*-Cre), astrocytes (*Gfap*-Cre), oligodendrocytes (*Olig2*-Cre) and microglia (*Cx3cr1*-Cre), respectively. (B) Immunoblot analyses of LysoIPs from neurons, astrocytes, oligodendrocytes, and microglia. LAMP1 and Cathepsin B were used as lysosomal markers, while Golgin-97, VDAC1, and Calreticulin served as markers for the Golgi, mitochondria, and endoplasmic reticulum, respectively. (C) LysoIP efficiency plot showing for whole brain and each cell type, including respective mockIP controls for each batch, the relative protein quantity of each replicate. Red barplots show the mean relative protein quantity of core lysosomal proteins found in each replicate, against the rest of the proteins quantified in gray. While in the negative control samples (mockIPs) the protein quantity between lysosomal proteins and the rest is similar and relatively low, the lysosomal protein content is enriched in all Cre positive samples, including the whole brain and the cell type specific Cre samples.

### The proteomics landscape of brain lysosomes

To characterize the proteomic landscape of brain lysosomes across different cell types, we first generated a list of proteins enriched in the LysoIP fraction relative to the mockIP controls. To avoid arbitrary cut-offs that could be influenced by IP efficiency or cell-type abundance, we applied a data-driven strategy based on the enrichment calculated from each Cre line, including whole-brain LysoIP (*CMV*-Cre)(Figure S2A). First, we calculated for each protein an enrichment score by comparing its levels in the LysoIP to that in the control mockIP dataset. Second, we applied a classifier based on well-defined core lysosomal proteins as the positive set (Table S1B), and nuclear chromatin-associated proteins as the negative set (Table S1C, see methods for details). This strategy validates the selective enrichment of lysosomal proteins by our approach, with an AUC (area under the curve) of 0.88-0.92, indicating high accuracy. It further enables the unbiased assignment of LysoIP-enriched proteins independently for each dataset (Figure S2B). Combining the enriched proteins (FPR < 0.05) from all Cre lines (Tables S2A-E), we identified 777 proteins that are LysoIP-enriched in at least one dataset (Figure 2A and Tables S2F). 170 out of the 777 proteins were found enriched in all datasets, and the majority of them (140/170, 82%) were previously annotated as lysosomal proteins (Figure 2A, Tables S1A and S2F). Gene ontology analysis^24^ confirmed significant enrichment for lysosome-related terms among LysoIP-enriched proteins, whereas even closely related terms like the endosome were enriched to a much lesser extent (e.g., lysosome term ORA score 4.77 vs. endosome ORA score 1.98)(Figures 2B, S2C and Tables S3A-C).

**Figure 2.**
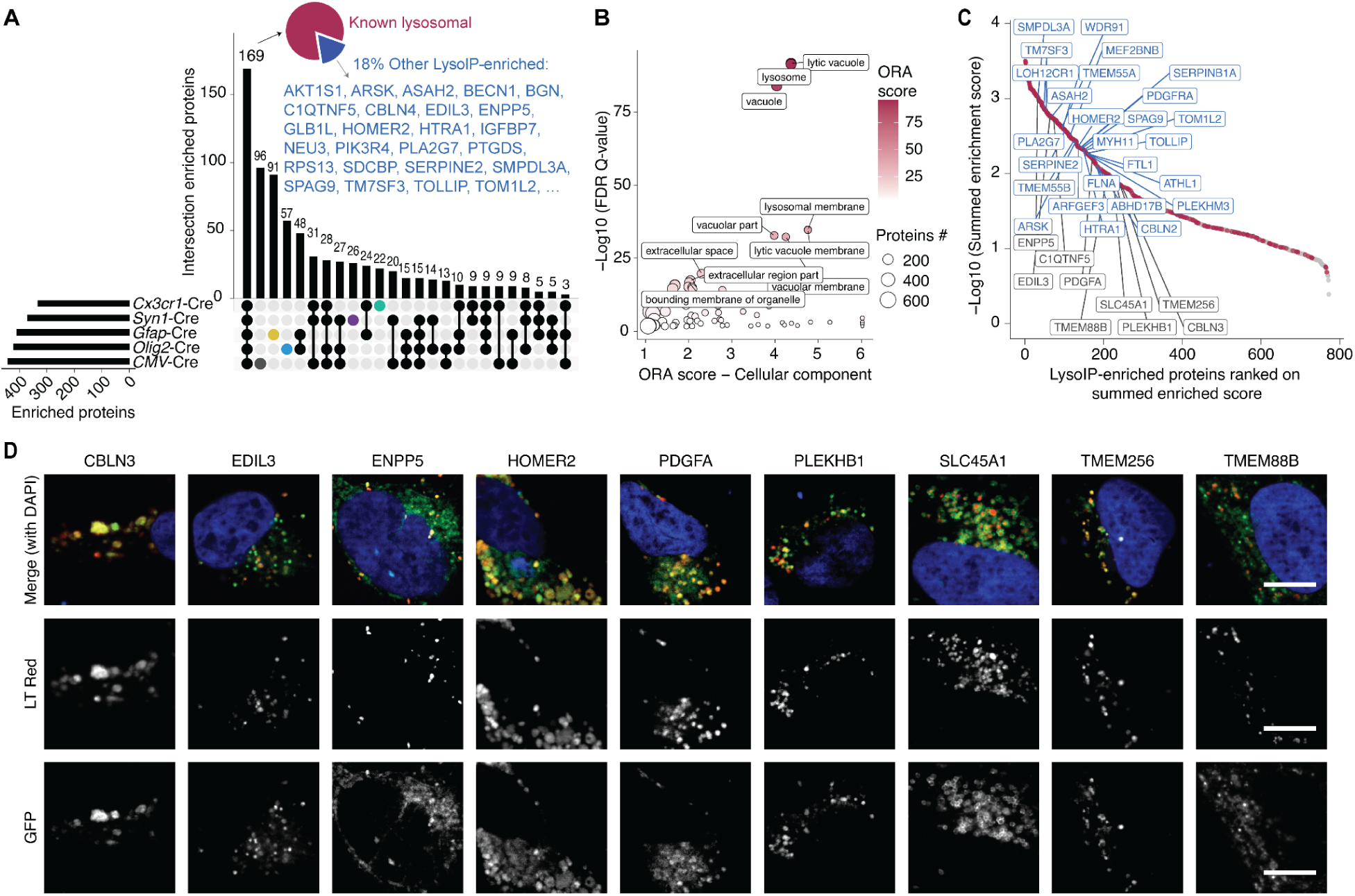
The proteomic landscape of brain lysosomes. (A) Upset plot showing overlap between LysoIP-enriched proteins in the different Cre lines (Tables S2A-E; *n* = 3-6). 82% of the proteins identified in all the Cre lines are known lysosomal proteins. Proteins highlighted in blue represent LysoIP-enriched proteins not yet reported as lysosomal. Known lysosomal proteins based on Table S1A. (B) Gene ontology (GO) terms over representation analysis (ORA) among LysoIP-enriched proteins. GO Cellular components terms are shown. ORA was performed using GOrilla (Gene Ontology enRIchment anaLysis and visuaLizAtion tool), with a *P*-value threshold of 0.001 using the list of all proteins detected and quantified in this study as background set. Top enriched terms according to ORA-score (GO enrichment combined with -log10(FDR *Q*-value)) are highlighted (see Table S3A). (C) LysoIP-enriched proteins were ranked based on their combined enrichment score across all Cre lines (see Table S2F). Red dots represent known lysosome-associated proteins based on Table S1A, gray dots represent other LysoIP-enriched proteins, top ranking selected proteins highlighted in blue. Proteins highlighted in gray boxes have been selected for validation by colocalization microscopy in Figure 2D. (D) Qualitative analysis of selected LysoIP-enriched proteins by fluorescence microscopy. Colocalization of transiently transfected GFP-tagged candidate proteins (GFP, green) and Lysotracker (LT, red) in U2OS cells. Nuclear staining in blue using DAPI. Scale bar is 10 µm.

**Figure S2.**
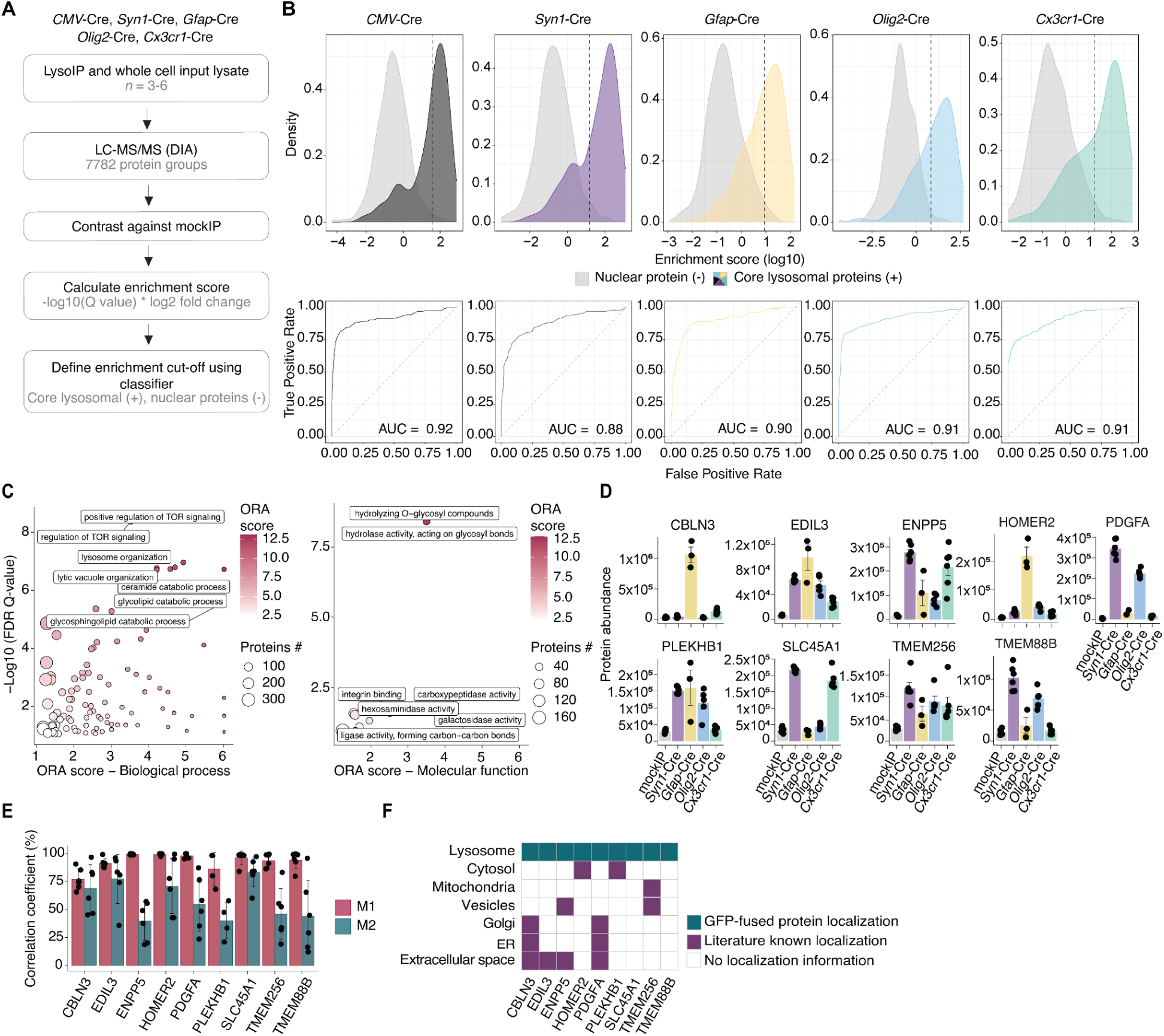
Identification of LysoIP-enriched proteins in brain cell types. (A) Workflow for analysis of proteomics data and defining LysoIP-enriched proteins. (B) Top panels: Plots showing classifiers for different Cre lines. The dashed lines indicate the enrichment score cut-off used for defining LysoIP-enriched proteins using a FPR < 0.05. Bottom panels: receiver operating characteristic (ROC) curves. The ROC curve illustrates the balance between sensitivity (true positives) and specificity (false positives), while the AUC (Area Under the Curve) provides a measure of overall performance of the classifier. A good classifier will have an ROC curve that hugs the top-left corner and an AUC close to 1. The dashed line represents a baseline for random performance. (C) Gene ontology (GO) terms over representation analysis (ORA) among LysoIP-enriched proteins. GO cellular compartments and molecular functions are shown. ORA was performed using GOrilla (Gene Ontology enRIchment anaLysis and visuaLizAtion tool, with a *P*-value threshold of 0.001 using the list of all proteins detected and quantified in this study as background set. Top enriched terms according to ORA-score (-log10(FDR q-value) plus GO enrichment) are highlighted (see also Tables S3B and S3C). (D) Barplot of protein abundance profiles of selected LysoIP-enriched proteins that were not previously reported as lysosomal proteins (highlighted in Figure 2C) (*n* = 3-6, error bars SEM). (E) Co-localization quantification by Manders Coefficient (MOC). M1 and M2 describe the contribution of each fluorescent channel from two selected channels to the pixels of interest. M1 being the fraction of LysoTracker overlapping with GFP signal (if M1 = 1: all lysosomes are GFP positive) and M2 is the fraction of GFP overlapping with LysoTracker (if M2 < 1: GFP also in other compartments than lysosome). Results are shown as median ± SD (*n* = 4-6 cells analyzed by Fiji with JaCoP-BIOP plugin, see methods). (F) All selected candidates showed to be localized to the lysosomal and also some to other subcellular compartments, as seen in Figure 2D. The localization of these proteins to other compartments as has been partially reported in the literature.

Among the brain LysoIP-enriched proteins, 302 were previously established to associate with the lysosome while 475 have not been so (Table S4), potentially representing new core and peripheral lysosomal proteins, or tissue-specific cargo destined for degradation. To validate the lysosomal localization, we ranked all LysoIP-enriched proteins based on their enrichment score across all Cre lines (Figure 2C, Table S2F), and selected 9 highly ranking proteins for experimental validation: CBLN3, EDIL3, ENPP5, HOMER2, PDGFA, PLEKHB1, SLC45A1, TMEM88B and TMEM256 (Figure S2D). We detected robust lysosomal localization in U2OS cells using GFP-tagged fusion of the nine candidates (Figures 2D and S2E). Of note, some of these proteins were reported to also localize to other compartments (Figures S2E, S2F), suggesting potential roles at inter-organelle contact sites^25,26^.

Altogether, the lysosomal protein atlas for brain cells provides a major resource for identifying novel lysosome-associated proteins.

### Cell-type-specific lysosomal proteomic landscape in brain cells

To determine the relative abundance of proteins in the lysosomes of different brain cell-types, we compared the normalized levels of proteins from LysoIP fractions of each Cre line. To this end, we normalized LysoIP-enriched protein abundances to the median levels of core lysosomal proteins, thereby controlling for cell number and LysoIP efficiency (Figure S3A). Normalized protein abundances, used *hereafter*, were highly correlated across cell types, indicating the overall similar lysosomal composition (Figure S3B). Consistently, the distribution of lysosomal proteins’ abundance across different metabolic functions was also overall similar across cell types (Figure S3C and S3D).

Although lysosomal composition was largely similar, principal component analysis (PCA) revealed distinct cell-type-specific lysosomal signatures, indicating unique proteomic landscape of lysosomes from different cell types (Figure 3A). Analysis of variance (ANOVA) across different cell types (Figure 3B and Table S4) revealed a subset of proteins whose lysosomal abundance is differential between brain cell types (Figures 3C and 3D). High ranking differential abundant proteins included well established cell type markers, e.g., CD68 for microglia, lysosomal membrane proteins, e.g., the proton-coupled amino acid transporter (PAT, SLC36A1) more abundant in neurons, as well as lumenal enzymes such as cathepsins and exonucleases, e.g., CTSH and PLD4 abundant in microglia, and sulfatases, e.g., the arylsulfatase G (ARSG) abundant in oligodendrocytes (Figures 3C and 3D).

**Figure 3.**
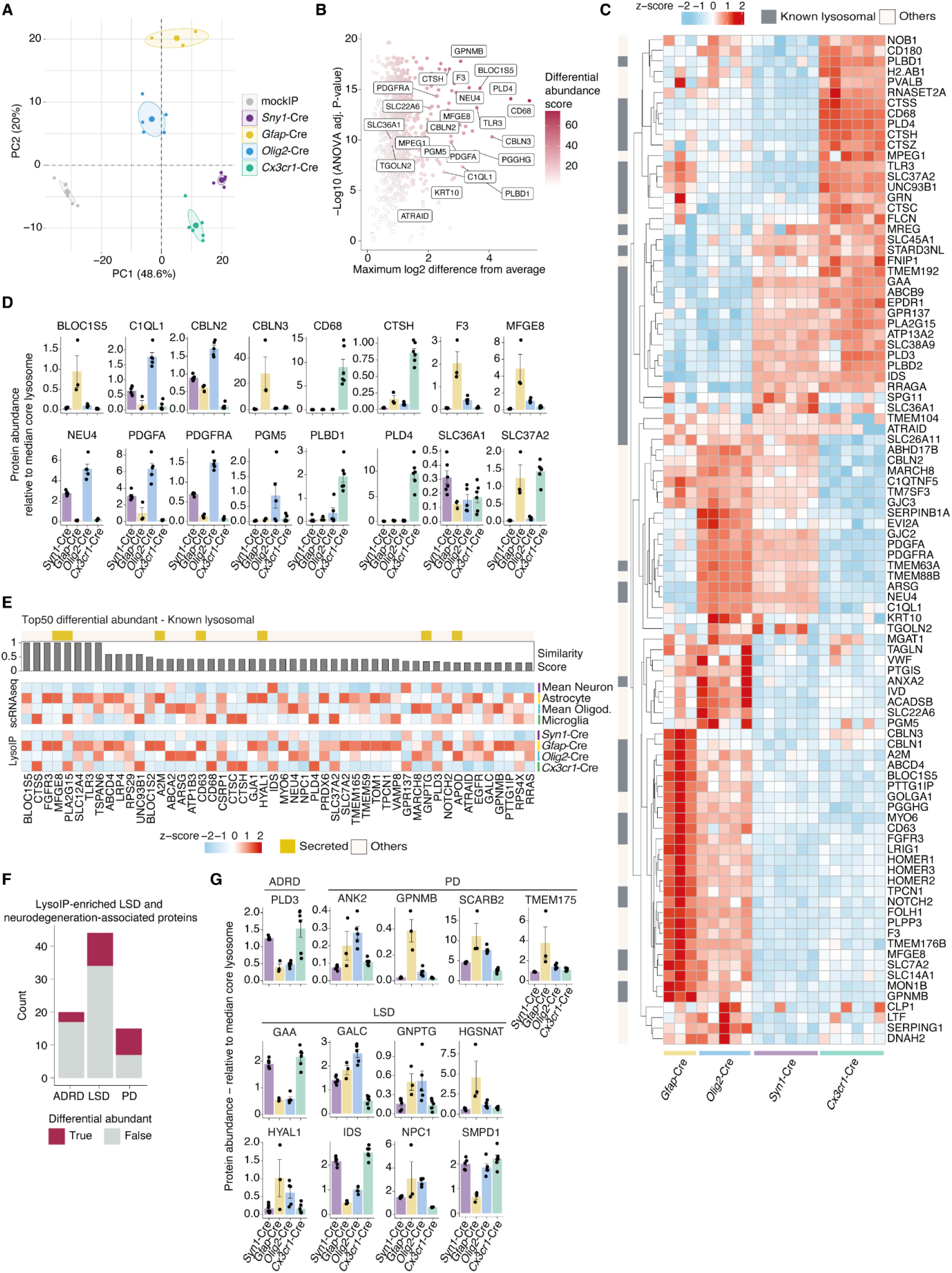
Cell-type-specific lysosomal proteomic landscape in the brain. (A) Principal component analysis (PCA) based on the normalized abundance of LysoIP-enriched proteins across Cre lines (*n* = 3-6; ellipses represent 95% confidence interval for each group). (B) Volcano plot showing differential protein abundance between LysoIP from different Cre lines. Log2-transformed protein abundances were normalized to the median of core lysosomal proteins and tested using ANOVA (Figure S3A, Table S4). Selected top differentially abundant proteins are highlighted. Proteins were ranked according to a differential abundance score calculated as -log10(ANOVA adjusted *P*-value) plus the maximum log2 difference from the mean abundance across all the Cre lines. (C) Heatmap of top 25 most differentially abundant proteins for each Cre line (80 proteins in total; *n* = 3-6; z-score transformed). Known lysosomal proteins (Table S1A) are highlighted on the left. (D) Barplots of top 6 most differentially abundant proteins per Cre line (16 proteins in total; ANOVA adjusted *P*-value < 0.05; *n* = 3-6; error bar SEM). (E) Heatmap comparison of protein abundance from LysoIP proteomics and mRNA levels for top differentially abundant proteins across brain cell types. mRNA levels were derived from a single cell RNAseq dataset obtained from DropViz^27^. The similarity between protein and mRNA abundance for each gene was calculated by computing the euclidean distance between ranks of abundances for mRNA and protein across cell types and using this distance to calculate a similarity score (see methods). RNAseq values for different neuron and oligodendrocyte subtypes were averaged (*n* = 3-6 for proteomic data; z-score transformed; Table S5). (F) Barplot of LSD- and neurodegeneration-associated proteins identified among LysoIP-enriched proteins. Differentially abundant proteins were defined with ANOVA adjusted *P*-value < 0.05 and absolute log2 abundance difference from the mean of all the Cre lines > 1. ADRD: Alzheimer’s disease and related dementia; PD: Parkinson’s disease; LSD: lysosomal storage disorders. (G) Barplot of differentially abundant LSD- and neurodegeneration-associated proteins (*n* = 3-6; error bar SEM).

**Figure S3.**
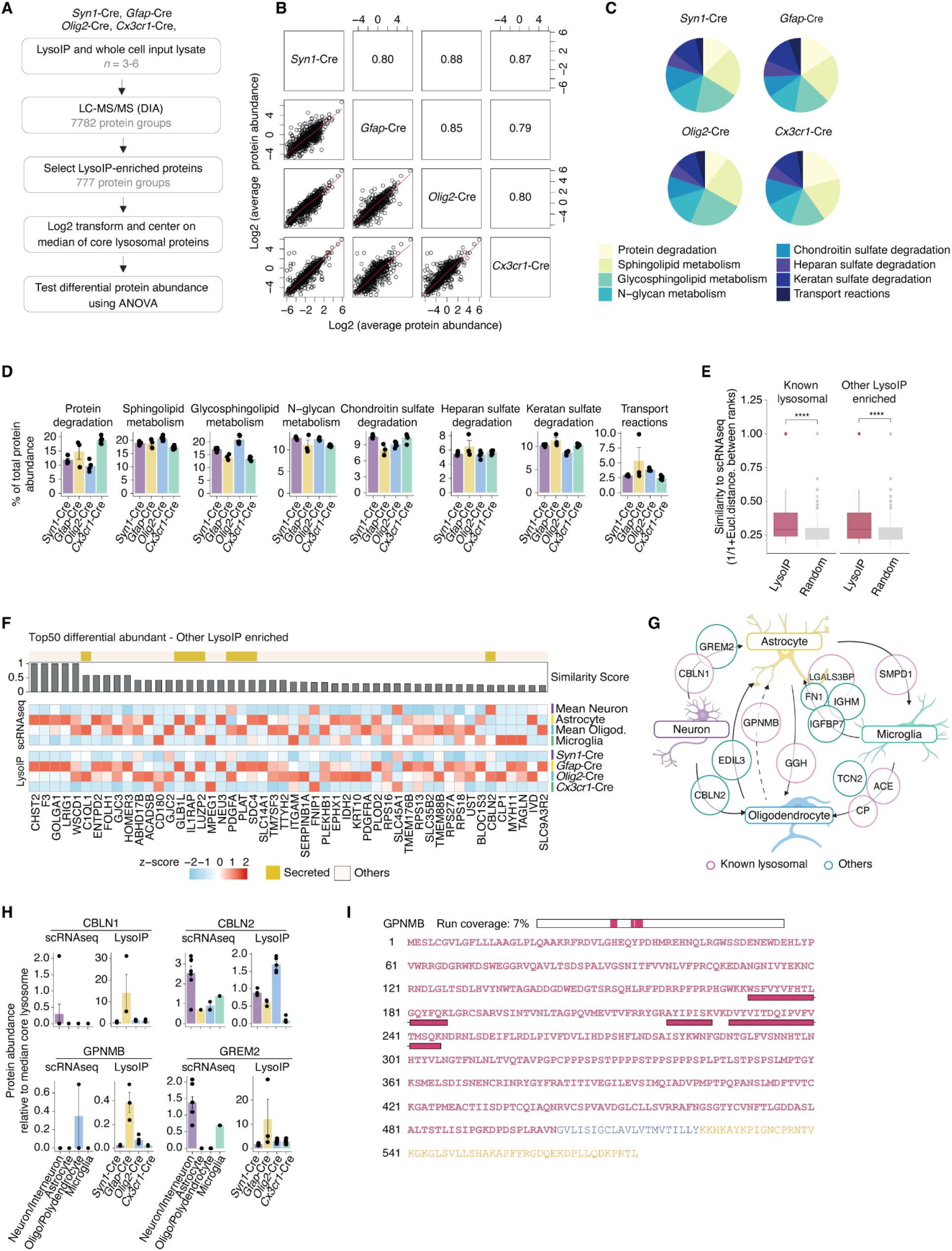
Differential protein abundance in LysoIPs from brain cell types. (A) Workflow to determine the differential abundance of LysoIP-enriched proteins between brain cell types. (B) Scatter plots showing the correlation of protein abundance across LysoIPs from different Cre lines. Log2-transformed protein abundances were normalized to the median of core lysosomal proteins and averaged between replicates (*n* = 3-6). The values indicated in the squares are Pearson correlation coefficients. (C) Pie chart of lysosomal metabolic functions across different cell types. Metabolic function annotations were derived from the metabolic atlas^33^. Normalized log2 protein abundances were summed for each metabolic function. (D) Barplots of the estimated percentage of total lysosomal protein abundance assigned to different metabolic functions (*n* = 3-6, error bars SEM). (E) Boxplot comparing similarity of cell-type specificity between single cell RNAseq and LysoIP proteome data. The two dataset exhibit higher levels of similarities across cell types compared to random (see methods) confirming isolation of cell-type-specific lysosomal proteins (Two-sample Wilcoxon test, RNAseq values for different neuron and oligodendrocyte subtypes were averaged; *n* = 3-6 for proteomic data)^27^, Table S5. (F) Heatmap comparison of normalized protein abundance from LysoIP proteomics and mRNA levels across brain cell types of selected proteins that were not previously reported as lysosomal proteins. Analysis as in Figure 3E. (G) Scheme of potential exchange of lysosomal proteins between brain cell types based on LysoIP data. Selected proteins displaying a low similarity score (< 0.24) between scRNAseq and LysoIP proteomics are shown (Table S5). The arrow starts from the cell type showing the highest mRNA expression for a given protein and points to the cell type with the highest protein abundance in LysoIP. (H) Barplots comparing mRNA expression from single cell RNAseq data and protein abundance from LysoIP proteomics for selected examples from panel G (RNAseq values for different neuron and oligodendrocyte subtypes were averaged; *n* = 3-6 for proteomics; error bar SEM)(Table S5). (I) Peptide coverage of the detected proteotypic peptides (pink bars), aligned to the primary sequence of GPNMB (UniProt accession number: Q99P91). The bar on top shows an overview map of the full sequence and its coverage. Highlighted in pink is the extracellular protein sequence, in blue the transmembrane region and in orange the cytoplasmic region (analyzed with Protter V1.0).

We next investigated potential mechanisms driving the differential abundance of lysosomal proteins across brain cell types. To test whether these differences could be explained by mRNA expression, we compared LysoIP proteomics to single-cell RNA sequencing data^27^. mRNA expression profiles were significantly correlated with the patterns of lysosomal protein abundance (“better than random”) for both known lysosomal and other LysoIP-enriched proteins (Figures 3E, S3E, S3F, and Table S5), thus validating our cell-type-specific LysoIP. Interestingly, several exceptions were observed, including secreted proteins like cerebellin 1 and 2 (CBLN1, CBLN2), gremlin 2 (GREM2), and the glycoprotein GPNMB, which were more abundant in lysosomes of non/less expressing cell types (Figure 3E). Three of these proteins are predominantly expressed by neurons, but are found to be more abundant in LysoIPs from astrocytes (CBLN1 and GREM2) and oligodendrocytes (CBLN2) (Figures 3E, S3G and S3H). GPNMB, however, is a transmembrane glycoprotein predominantly expressed by oligodendrocytes and activated microglia associated with disease^28^, and we found it to be highly abundant in astrocyte-derived lysosomes (Figures S3G and S3H), likely because of its shedding from the cell surface^29^. Detailed analysis of peptides identified for GPNMB confirmed their origin from the shed extracellular domain of the protein (Figure S3I). These results indicate that cell-type specific LysoIP enables detection of proteins produced in one cell type and uptaken/used in others (Figure S3G), an observation of major therapeutic importance for cell based therapies that depends on the principle of cross-correction^30,31^. Together these data show that the protein composition of lysosomes varies between brain cell types, at least in part due to differences in the expression of genes encoding lysosomal proteins, and that cell-type specific LysoIP can be used to uncover novel lysosomal protein dynamics potentially mediating inter-cellular communication.

Mutations in lysosomal genes are major risk factors for neurological diseases^5,10,11^, to investigate if proteins encoded by these genes are differentially abundant in lysosomes of different cell types, we analyzed the enrichment patterns for a set of genes that have been genetically linked to LSDs, Parkinson’s Disease (PD), and Alzheimer’s Disease Related Dementias (ADRD). Across these three disease groups, we detected 67 LysoIP-enriched proteins, the majority of which are present at comparable abundances in lysosomes from brain cell types (Figure 3F and Table S4). Interestingly, we found a subset of disease-associated lysosomal proteins to be differentially abundant in LysoIPs from different Cre lines, suggesting a potential cell-type-specific role of disease-associated proteins (Figure 3G). We have also identified an additional set of LysoIP-enriched proteins that occur in the online mendelian inheritance in man (OMIM) database^32^, including a subset of potentially newly discovered lysosomal proteins (Table S4). Mutations in genes encoding these proteins may underlie previously unrecognized lysosomal storage disorders, offering new avenues and great resource for investigating lysosomal dysfunction in neurological diseases.

### SLC45A1 is a novel lysosomal protein expressed in neuronal cells

Among the disease-relevant proteins displaying differential abundance between brain cell types, SLC45A1, a transmembrane protein, caught our attention. Mutations in SLC45A1 cause a monogenic neurological disorder with neuropsychiatric features^14–16^, and was never reported as a lysosomal protein but suggested to be a neuron-specific plasma membrane sugar transporter^14,34^. We hypothesized that SLC45A1 is a novel neuron-specific lysosomal protein, and thus SLC45A1-associated disease is a previously unknown lysosomal storage disorder. Indeed, SLC45A1 is selectively expressed in the brain (Figure S4A)^34^ and its mRNA expression in primary brain cells is restricted to neurons (Figure S4B). We detected SLC45A1 in microglia-derived lysosomes, likely due to *Cx3cr1*-Cre leakage into neurons or neuronal pruning, as SLC45A1 mRNA was low or undetectable in microglia (Figures S2D, S4B and S4C). SLC45A1 expression increases upon differentiating human iPSCs to neurons (iNeuron) both at mRNA (Figure S4D) and protein levels using endogenously tagged SLC45A1 protein (Figures 4A and 4B). Furthermore, we observed that SLC45A1 is expressed in human neuroblastoma SH-SY5Y cells (Figure S4D) and its expression increased under serum deprivation (Figure S4E), a condition used in our functional analysis to increase the dependence on SLC45A1 in these cells.

**Figure 4.**
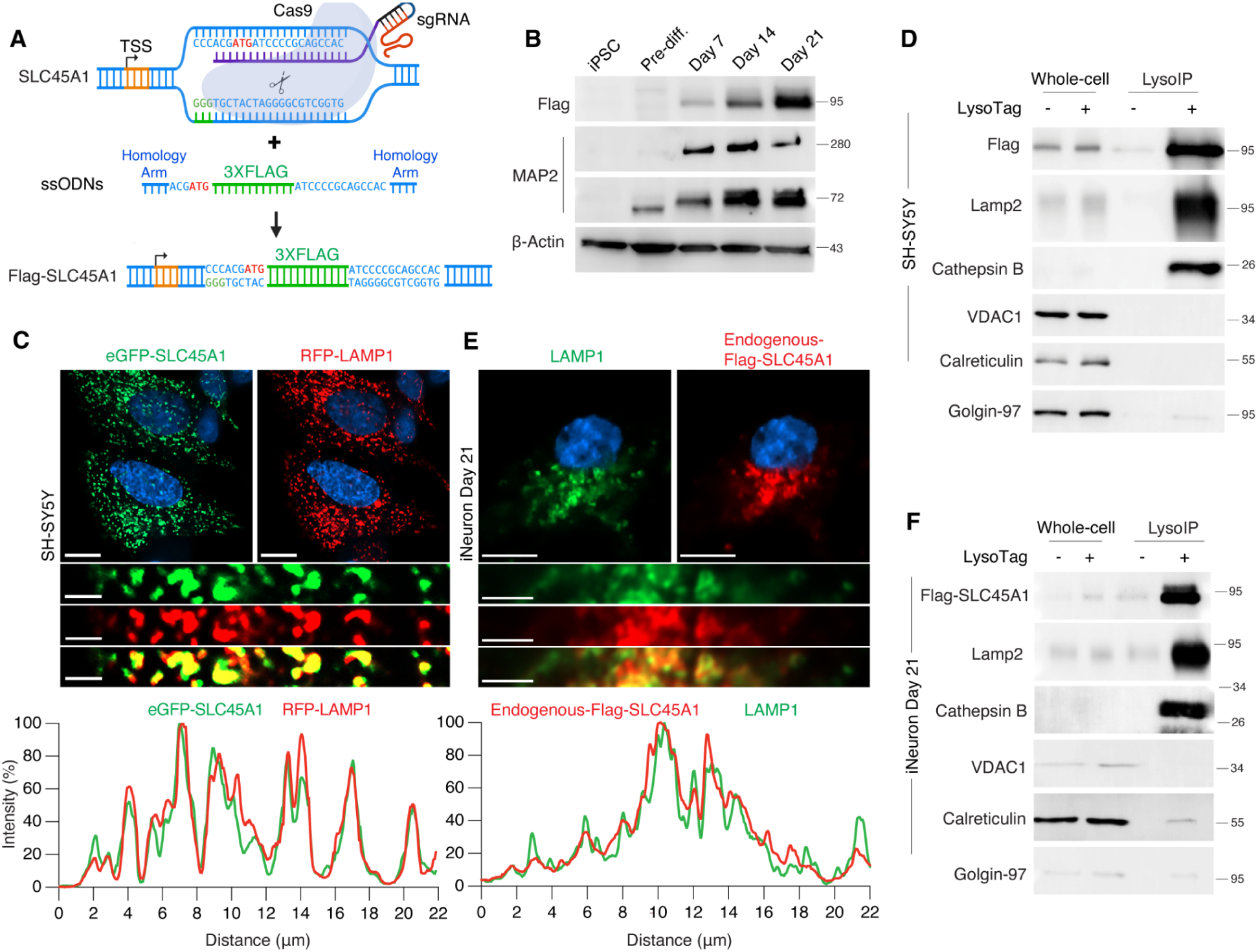
SLC45A1 is a lysosomal protein. (A) Schematic representation of the endogenous Flag-tagging of SLC45A1 using CRISPR/Cas9 technology in iPSCs. ssODNs: single-stranded oligodeoxynucleotides. (B) Immunoblot analyses of endogenous Flag-tagged SLC45A1 expression in iPSCs and at pre-differentiation (pref-diff.) stages, as well as on days 7, 14, and 21 post-neuronal differentiation. A Flag antibody was used to detect Flag-SLC45A1 fusion, and MAP2 served as a neuronal differentiation marker. (C) Live imaging to assess the co-localization of eGFP-SLC45A1 (green) with RFP-LAMP1 (red) in SH-SY5Y cells. Representative image is shown. Scale bars are 10 µm (top) and 2 µm (bottom). A representative pixel intensity plot is shown for RFP-LAMP1 (red) and eGFP-SLC45A1 (green). (D) Immunoblot analyses to detect Flag-tagged SLC45A1 in the lysosomal fraction (LysoIP) of SH-SY5Y cells. LAMP2 and Cathepsin B were used as lysosomal markers, while Golgin-97, VDAC1, and Calreticulin were used as markers for the Golgi, mitochondria, and endoplasmic reticulum, respectively. (E) Immunofluorescence analysis to assess the co-localization of endogenous Flag-tagged SLC45A1 in iPSC-derived neurons (day 21) and the lysosomes. LAMP1 was used as a lysosomal marker (green), and a Flag antibody was used to detect endogenous SLC45A1 (red). Representative image is shown. Scale bars are 5 µm (top) and 2 µm (bottom). A pixel intensity plot is shown for LAMP1 (green) and Flag-SLC45A1 (red). (F) Immunoblot analyses to detect endogenously Flag-tagged SLC45A1 in the lysosomal fraction (LysoIP) of iNeurons (day 21). LAMP1 and Cathepsin B were used as lysosomal markers, while Golgin-97, VDAC1, and Calreticulin were used as markers for the Golgi, mitochondria, and endoplasmic reticulum, respectively.

**Figure S4.**
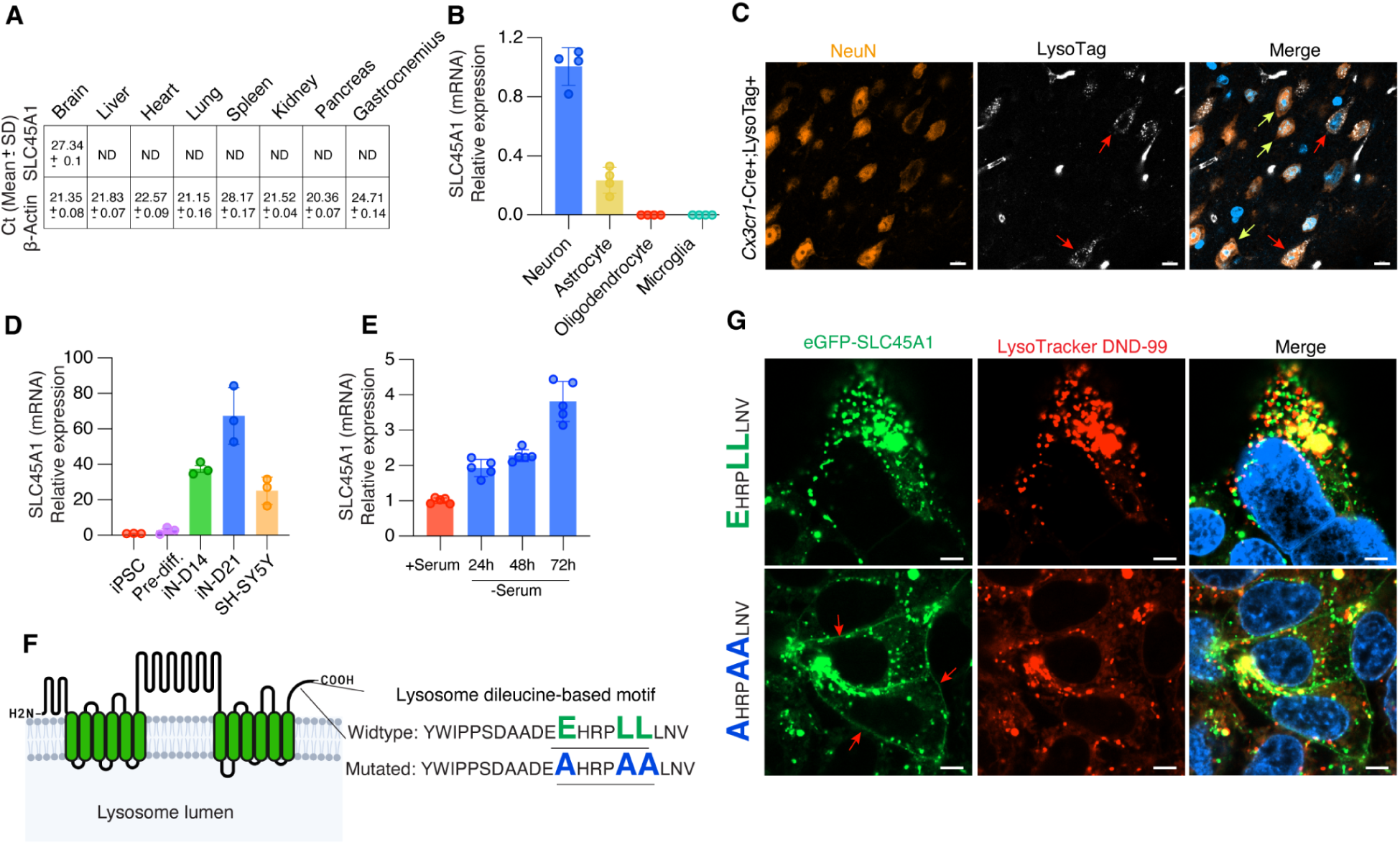
Characterization of SLC45A1 expression. (A) Quantitative RT-PCR analysis of SLC45A1 mRNA across mouse tissues. The cycle threshold (Ct) value indicates the number of amplification cycles required for SLC45A1 mRNA to exceed the basal threshold level. ND represents samples where Ct was not detected. One of the replicates of kidney and pancreas was above 37, which was considered ND as well. Representative experiment with *n* = 3 technical replicates. β-Actin was used as endogenous control. (B) Quantitative RT-PCR analysis of SLC45A1 mRNA expression in different mouse brain primary cell types, including neurons, astrocytes, oligodendrocytes, and microglia. β-Actin was used as an endogenous control to normalize gene expression levels. Relative mRNA abundance in each cell type compared to neurons is reported. Results represent mean ± SD (Representative experiment with *n* = 4 technical replicates). (C) Immunofluorescence analysis to assess the co-localization of the LysoTag with NeuN, a neuronal marker, in *Cx3cr1*-Cre+;LysoTag+ mouse brain sections. Red arrows indicate neurons expressing LysoTag, while yellow arrows highlight neurons that do not express LysoTag. Scale bar is 10 µm. (D) Quantitative RT-PCR analysis of human SLC45A1 mRNA expression in induced pluripotent stem cells (iPSCs), cells at pre-differentiation stages, and cells at 14 and 21 days post-differentiation into induced neurons (iN), as well as in SH-SY5Y cells. β-Actin was used as the endogenous control for normalization. Relative mRNA abundance in each cell type compared to iPSCs is reported. Results represent mean ± SD (Representative experiment with *n* = 3 technical replicates). (E) Quantitative RT-PCR analysis of human SLC45A1 mRNA expression in SH-SY5Y cells under serum starvation conditions. β-Actin was used as the endogenous control for normalization. Relative mRNA abundance in each cell type compared to +serum condition is reported. Results represent mean ± SD (Representative experiment with *n* = 5 technical replicates). (F) Topology prediction of SLC45A1 at the lysosomal membrane analyzed by Protter V1.0 with identification of lysosomal dileucine motif: EXXXLL at the C-terminus. Our mutagenesis strategy to AXXXAA is shown. (G) Live cell imaging to assess the co-localization of wildtype or mutated FLAG-eGFP-SLC45A1 (green) with LysoTracker DND-99 (red) as lysosomal marker in HEK293T cells. Scale bar is 5 µm.

We then rigorously validated the lysosomal localization of SLC45A1. First, we established its strict lysosomal localization, using immunofluorescence, when exogenously expressed in U2OS (Figure 2D), HEK293T cells (Figure S4G) and in SH-SY5Y cells (Figures 4C). Of importance, we identified a consensus acidic dileucine-based motif, EXXXLL, at the C-terminus of the SLC45A1 protein (Figures S4F and S4G) whose mutagensis to AXXXAA resulted in a fraction of the protein to mislocalize to the plasma membrane although a large fraction remained on the lysosome (Figure S4G). In line with the microscopy data, LysoIP confirmed lysosomal localization of SLC45A1 in SH-SY5Y cells (Figure 4D). Finally, endogenously tagged SLC45A1 (Figures 4A and 4B) localized entirely to the lysosome in iNeurons, as confirmed by both microscopy and LysoIP (Figures 4E and 4F), ruling out over-expression artifact and establishing SLC45A1 as a core lysosomal protein in neuronal cells, thus underscoring its potential role in lysosomal function.

### SLC45A1 loss leads to lysosomal dysfunction

To investigate whether SLC45A1 loss causes a previously undescribed lysosomal storage disorder, we generated and validated SLC45A1-deficient SH-SY5Y cells (Figures S5A, S5B, Table S6). Consistent with cellular hallmarks of LSDs^35,36^, SLC45A1 loss led to a significant increase in LAMP1 signal and perinuclear lysosomal clustering (Figure 5A and S5C). Transmission electron microscopy (TEM) confirmed an increase in lysosomal size. It also revealed the accumulation of dense lysosomes with lipofuscin-like pigments, without an increase in lysosomal numbers (Figure 5B). Furthermore, SLC45A1-deficient cells showed a substantial blockade in autophagic flux when compared to wildtype cells (Figure 5C), and their lysosomes exhibited a significant reduction in proteolytic capacity (Figure S5D). Importantly, all these phenotypes were reversed by re-expressing SLC45A1 (Figures 5A-C, S5C and S5D). Notably, untargeted lipidomics revealed a significant increase in bis(monoacylglycero)phosphate (BMP) levels, lysosomal lipids known to be elevated in numerous LSDs (Figure 5D and E)^37^. Altogether, our data provide strong evidence for the presence of lysosomal dysfunction in SLC45A1-deficient cells, a hallmark of bona fide LSD.

**Figure 5.**
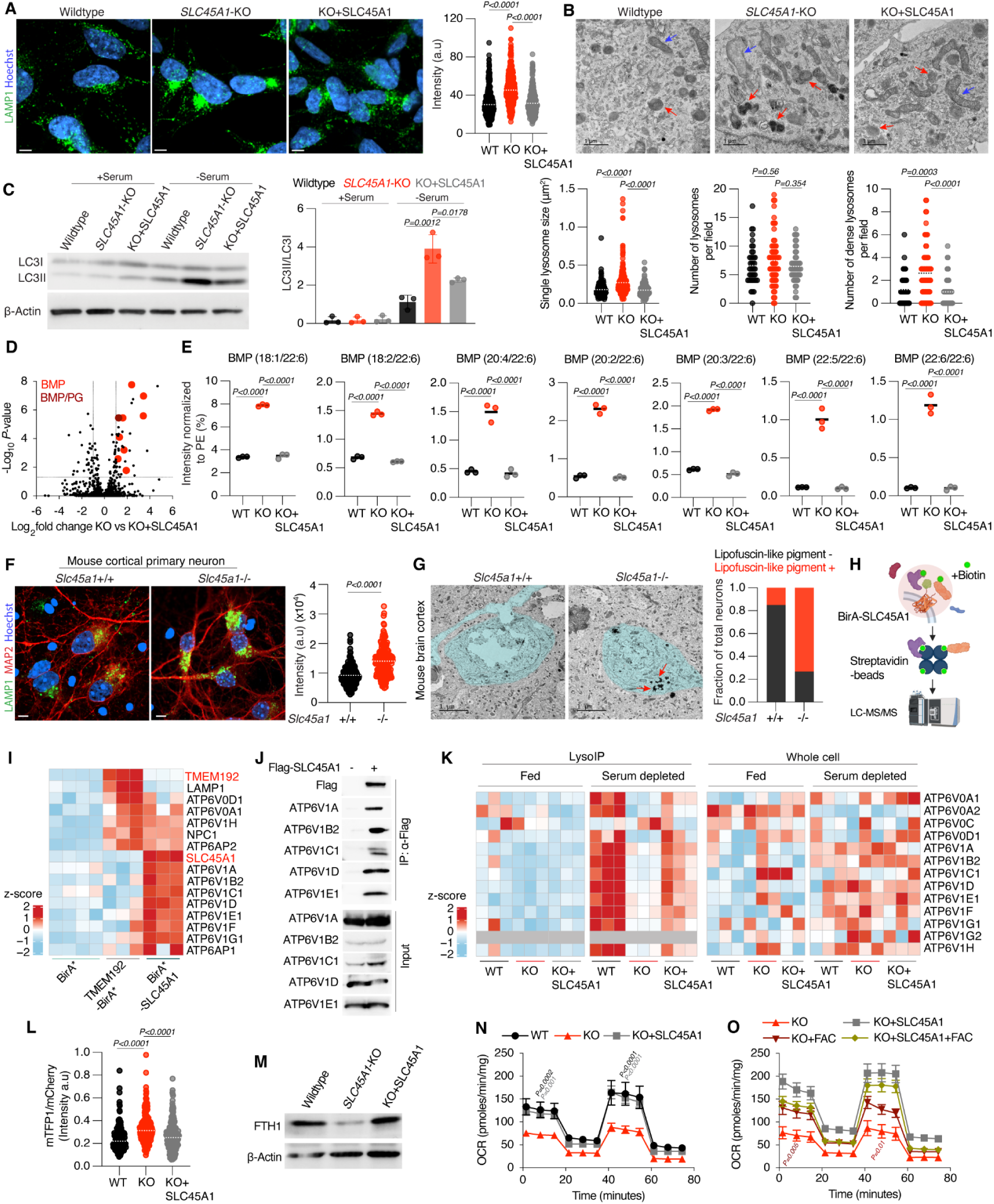
Loss of SLC45A1 leads to lysosomal dysfunction and mitochondrial impairment. (A) Immunofluorescence analysis of LAMP1 signal in human SH-SY5Y cells: wildtype (*n* = 20 fields), *SLC45A1*-KO (*n* = 26 fields), and rescued (KO+SLC45A1) (*n* = 23 fields). Representative images are included. Scale bar is 5 µm. Quantification on the right. *P*-values were calculated by one-way analysis of variance (ANOVA) with Tukey’s honestly significant difference (HSD) test. (B) Transmission electron microscopy (TEM) analysis of human SH-SY5Y. Representative images are shown. Red and blue arrows indicate lysosomes and mitochondria, respectively. Bottom panels show dot plots for single lysosome size (µm²), the number of lysosomes per field (5.54 µm x 5.54 µm), and the number of dense lysosomes per field (5.54 µm x 5.54 µm) in wildtype (*n =* 40 fields), *SLC45A1*-KO (*n* = 65 fields), and rescued cells (*n* = 50 fields). Horizontal dotted lines represent the mean. *P*-values were calculated by ANOVA with Tukey’s HSD test. (C) Immunoblot analysis of LC3I and LC3II in human SH-SY5Y cells: wildtype, *SLC45A1*-KO, and rescued (KO+SLC45A1) under serum replete (+serum) and deplete (-serum) conditions. Quantitation in the right panel. *P*-values were calculated by one-way ANOVA with Tukey’s HSD test. (*n* = 3, independent biological samples). (D) Untargeted lipidomics analysis of *SLC45A1*-KO and rescued (KO+SLC45A1) SH-SY5Y cells under 72 h serum deprivation conditions. Data is presented as a volcano plot of log_2_-transformed fold changes in the abundance of lipids between *SLC45A1*-KO and rescued cells. The horizontal line indicates a *P*-value of 0.05 and the vertical lines indicate a fold change of 2. *n* = 3 biological independent samples per genotype. *P*-values were calculated by one-way analysis of variance (ANOVA) with Tukey’s honestly significant difference (HSD) test. Lipidomics data for wildtype, *SLC45A1*-KO, and rescued cells are provided in Table S7. BMPs were identified with definitive -H and +NH_4_ MS2s. BMP/PG was used in cases where +NH_4_ MS2 data was not acquired for definitive distinction between isomeric BMP and PG. (E) Targeted quantitation of BMPs after normalizing to endogenous lipid PE(16:0/18:1) (POPE) in wildtype, *SLC45A1*-KO, and rescued cells (*n* = 3 each). *P*-values were calculated by one-way ANOVA with Tukey’s HSD test. (F) Immunofluorescence analysis and quantification of LAMP1 signal in primary neurons isolated from *Slc45a1*+/+ (*n =* 33 fields) and *Slc45a1*-/- mice (*n =* 31 fields)*. P*-values were calculated by two-tailed independent t-tests. Horizontal dotted lines represent the mean. Scale bar is 5 µm. (G) TEM analysis of mouse cortex from *Slc45a1*+/+ and *Slc45a1*-/- mice. Blue shadow indicates neuronal cells. Arrows (red) indicate lipofuscin-like pigment in the neurons. (H) Schematic overview of proximity labeling (BioID) experiment. BirA* (*n* = 4), and TMEM192-BirA* (*n* = 3) were used as controls. BirA*-SLC45A1 (*n* = 3). (I) Heatmap representation of selected lysosomal proteins enriched by BirA*-SLC45A1 or TMEM192-BirA* in the BioID experiment. Candidates were selected for being enriched over BirA* (negative control) with a mean log2 ratio > 1 and *Q*-value < 0.05. Since we found multiple members of the vATPase to be enriched in BirA*-SLC45A1, we included all the detected subunits in the heatmap. Some shared candidates are also included. Candidates selected based on Table S8A. Heatmap data based on Table S8B. *n* = 3-4 biologically independent samples. (J) Immunoblot analyses of anti-Flag immunoprecipitation of Flag-SLC45A1 in SH-SY5Y cells followed by detection of interacting endogenous V-ATPase subunits including V1A, V1B2, V1C1, V1D and V1E1. (K) Heatmap representation of relative protein abundance of V-ATPase subunits in LysoIP and whole-cell fractions from human SH-SY5Y wildtype, and *SLC45A1*-KO, and rescued cells under serum fed and starved conditions. Data provided in Table S9A-B. *n* = 3 biologically independent samples, *n* = 2 for whole cell KO +SLC45A1 fed condition. Significantly depleted V-ATPase subunits in KO vs WT are indicated. Protein complex selected based on BioID experiment in (I), that was found to interact with SLC45A1. Gray boxes indicate that proteins were not detected. (L) Assessment of lysosomal acidity using the FIRE-pHLy reporter (see S5J) in SH-SY5Y cells. Fluorescence intensity ratios for mTFP1 and mCherry (mTFP1/mCherry) were calculated. Increased ratio indicates increase in lysosomal pH. *P*-values were calculated by one-way ANOVA with Tukey’s HSD test. Horizontal dotted lines represent the mean. (M) Immunoblot analysis of ferritin heavy chain (FTH1) with β-Actin as loading control in whole-cell lysates prepared from wildtype, *SLC45A1*-KO and rescued cells upon serum deprivation. (N) Mitochondrial oxygen consumption rates (OCR) in wildtype (*n* = 4), *SLC45A1*-KO (*n* = 6) and rescued SH-SY5Y cells (*n* = 4). Reads were normalized to total protein. Results represent mean ± SEM. *P*-values were calculated by two-way ANOVA test and reported against the KO with the color matches that to which it is compared. (O) Mitochondrial OCR upon iron supplementation with 0.2 mg/ml ferric ammonium citrate (FAC) in *SLC45A1*-KO and rescued SH-SY5Y cells. Reads were normalized to total protein. Results represent mean ± SEM (*n* = 3). *P*-values were calculated by two-way ANOVA test and reported against the KO with the color matches that to which it is compared.

**Figure S5.**
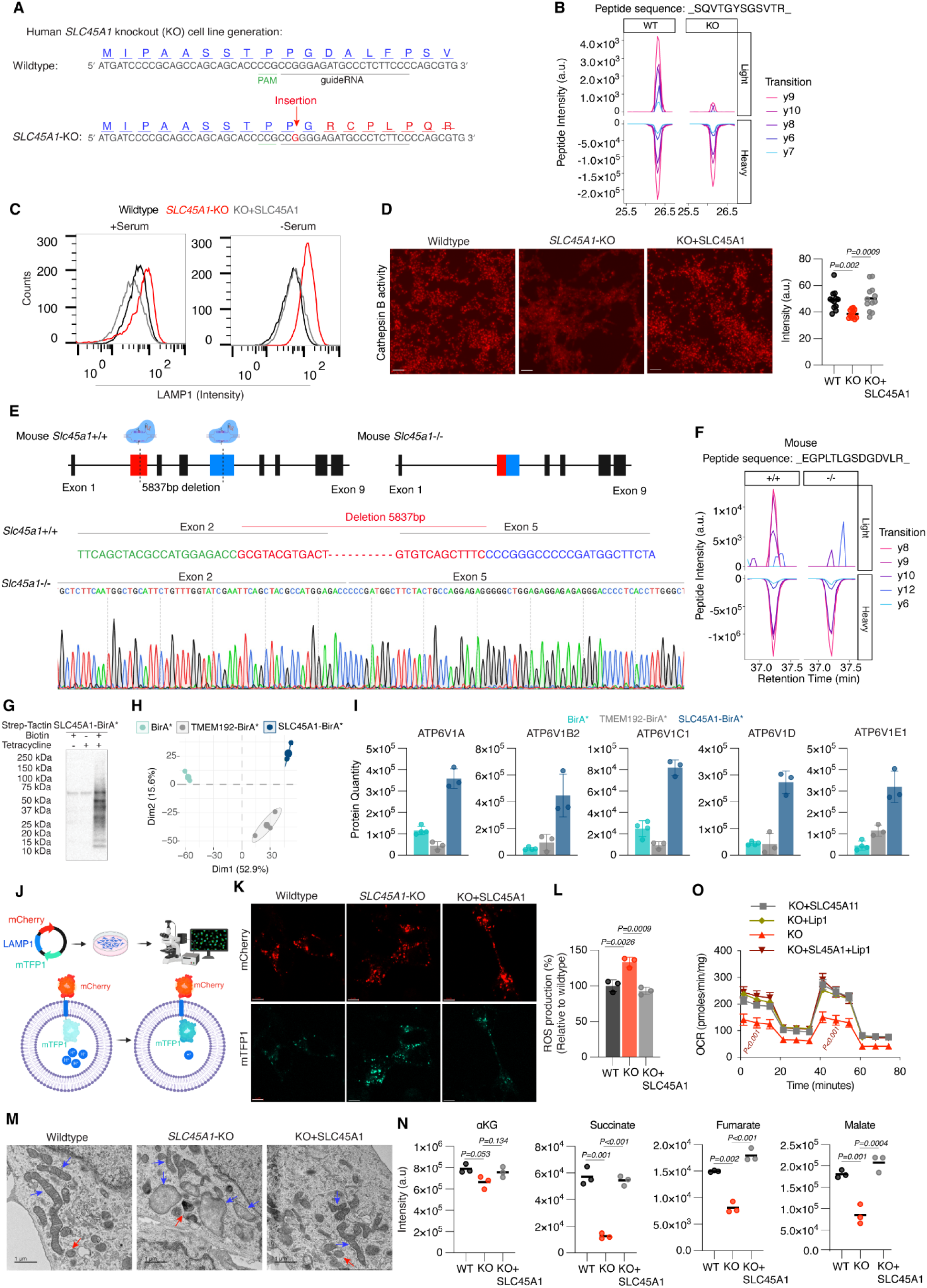
Generation and characterization of SLC45A1-deficient human SH-SY5Y cells and mice. (A) Successful generation of SH-SY5Y *SLC45A1*-KO cells using CRISPR/Cas9 as evident from sequencing results showing a frameshift mutation caused by the addition of a guanosine nucleotide. sgRNA is underlined. Analysis was performed using Synthego ICE indel deconvolution tool. (B) Validation of SLC45A1 loss in *SLC45A1*-KO cells using parallel reaction monitoring (PRM) to detect SLC45A1 peptides in SH-SY5Y wildtype and KO cells. PRM uses heavy spike-in peptides. Co-elution of endogenous (light) and standard (heavy) peptides enables quantitation relative to the spike-in standard measured by LC-MS. (C) LAMP1 intensity, using flow cytometry, in wildtype, *SLC45A1*-KO, and rescued cells under serum replete and deplete (for 72 h) conditions. (D) Live-cell imaging to assess Cathepsin B activity in wildtype, *SLC45A1*-KO, and rescued cells after under serum deprivation (72 h) using the Magic Red Cathepsin B activity assay. Quantification to the right. *P*-values were calculated using one-way ANOVA with Tukey’s HSD test (*n* = 12 fields each). (E) Schematic overview of the strategy to generate *Slc45a1* knockout mice. Guide RNAs were designed to target exon 2 and exon 5, resulting in a deletion of 5837 bps. (F) Validation of SLC45A1 loss at peptide level in *Slc45a1*-/- (KO) mice using PRM as in panel B. (G) Immunoblot validation of biotinylation efficiency of BirA*-SLC45A1 of the BioID experiment. Expression measured in whole-cell lysates collected from HEK293 cells, following 24 h incubation with or without (+/-) 1 µg/ml Tetracycline and (+/-) Biotin. (H) Principal component analysis (PCA) based on the relative abundance of BioID-enriched proteins between BirA* constructs (*n* = 3; ellipses represent 95% confidence interval for each group). (I) Barplots showing relative protein quantity of selected V-ATPase subunits between controls (BirA* and TMEM192-BirA*) and BirA*-SLC45A1 (*n* = 3; Table S8B). (J) Schematics of lysosomal acidity assessment protocol. FIRE-pHLy is targeted to lysosomes via LAMP1. The lysosomal pH-sensitive mTFP1 (cyan fluorescent protein) is located in the lumen, and the lysosomal pH-insensitive mCherry (red) is positioned on the cytosolic side. To assess lysosomal acidity, fluorescence intensity ratios of mTFP1 to mCherry (mTFP1/mCherry) were calculated (Figure 5L). (K) Representative individual channel images of FIRE-pHLy-expressing SH-SY5Y wildtype, *SLC45A1*-KO, and rescued cells. Scale bar is 5 µm. (L) Measurement of reactive oxygen species (ROS) using CellROX in SH-SY5Y wildtype, *SLC45A1*-KO, and rescued cells after 72 h of serum deprivation using flow cytometry. Relative change (%) compared to wildtype is reported. *P*-values were calculated by one-way ANOVA with Tukey’s HSD test (*n* = 3). (M) Representative TEM images of SH-SY5Y wildtype, *SLC45A1*-KO, and rescued cells. Blue arrows indicate mitochondria and red arrows show lysosomes. Scale bar is 1 µm. (N) Targeted quantitation of alpha-ketoglutarate (ɑKG), succinate, fumarate, and malate in wildtype (WT), *SLC45A1*-KO, and rescued SH-SY5Y cells from the experiment in Figure 6A. *P*-values were calculated using one-way ANOVA with Tukey’s HSD test (*n* = 3). (O) Mitochondrial oxygen consumption rates (OCR) in *SLC45A1*-KO (*n* = 4), rescued cells (*n* = 4) upon 250 nM liproxstatin-1 (Lip1) (*n* = 3) treatment. Reads were normalized to total protein. Results represent mean ± SEM. *P*-values were calculated by two-way ANOVA test.

To assess whether SLC45A1 loss causes lysosomal dysfunction in vivo, we generated a *Slc45a1* knockout mouse with a 5.8 kb deletion spanning exons 2 to 5 (Figure S5E), confirmed by PRM mass spectrometry (Figure S5F). Consistent with our cell line data, loss of SLC45A1 led to a significant increase in LAMP1 signal and its perinuclear clustering in primary cortical neurons (Figure 5F). TEM revealed a significant abundance of dense lysosomes with lipofuscin-like pigment in cortical neurons of the *Slc45a1* knockout mouse (Figure 5G), providing definitive evidence for lysosomal involvement in SLC45A1 deficiency *in vivo*.

SLC45A1 exhibits a topology with extensive cytosolic loops (Figure S4F), suggesting a structural role at the lysosomal surface, similar to other lysosomal transmembrane proteins^3^. To uncover such potential function and the impact of its loss on the lysosome, we used proximity labeling to identify interactors using biotinylation identification (BioID) (Figures 5H, S5G, and S5H). Strikingly, BirA*-SLC45A1, but not BirA* or TMEM192-BirA* controls, recovered more of five V1 subunits of the vacuolar ATPase (V-ATPase) complex including ATP6V1A, ATP6V1D, ATP6V1C1, ATP6V1E1, and ATP6V1B2 (Figure 5I and S5I). Co-immunoprecipitation confirmed the interaction between SLC45A1 and the V1 subunits (Figure 5J). Based on these results, we hypothesized that SLC45A1 stabilizes/recruits the V1 complex to the lysosome, thus regulating the lysosomal pH homeostasis. Indeed, we observed a remarkable decrease in the lysosomal abundance of V1 subunits in SLC45A1-deficient cells compared to that of the wildtype with no decrease in their whole-cell abundance (Figure 5K). Of importance, this phenotype is observed under starvation condition, in which the assembly of V1 subunit on the lysosome is enhanced^38–40^ (Figure 5K). Consistent with this depletion, we observed a significant increase in lysosomal pH in *SLC45A1*-KO cells compared to wildtype cells that is completely rescued by re-expressing SLC45A1 (Figures 5L, S5J and S5K). Impaired lysosomal acidification leads to cellular iron deficiency, increase in reactive oxygen species (ROS) and subsequent mitochondrial dysfunction in cell lines and mouse models of LSDs^41–43^. Indeed, we observed lower ferritin (Figure 5M) and higher ROS (Figure S5L) levels in *SLC45A1*-KO cells. Furthermore, loss of SLC45A1 leads to lower oxygen consumption rate (OCR) (Figure 5N) and alterations in mitochondrial morphology with the formation of vacuole-like structures^44^ (Figure S5M), consistent with mitochondrial dysfunction. Indeed, metabolite profiling showed that mitochondrial metabolites whose generation is dependent on iron-sulfur cluster-containing complexes (succinate, fumarate and malate)^43^ are reduced upon SLC45A1 loss (Figures S5N and 6A). Of importance, iron supplementation or ROS inhibitor (Liproxstatin-1) rescued the impaired mitochondrial OCR in SLC45A1-deficient cells (Figures 5O and S5O). Altogether, our results demonstrate that SLC45A1 loss induces lysosomal dysfunction consistent with LSD and further implicate a pathological mechanism through mitochondrial dysfunction in SLC45A1-associated disease.

### SLC45A1 is required for the efflux of hexose from the lysosome

Previous study showed that rat SLC45A1 contains several conserved motifs found in sugar transporters, including the RXGRR, which is also present in the human and mouse protein (Figure S6A)^34^. Indeed, multiple sequence alignment between the human, rat, and mouse SLC45A1 indicates a highly conserved sequence (∼90 to 97%) (Figure S6B). Untargeted metabolite profiling revealed a significant accumulation of hexose (C_6_H_12_O_6_) in SLC45A1-deficient cells (Figures 6A and 6B). These results are contrary to its proposed function as plasma membrane sugar importer^34^, which if correct predicts lower hexose levels, and consistent with potential sugar transport function in the lysosome. Thus, we hypothesized that SLC45A1 functions as a hexose sugar exporter at the lysosomal membrane. To directly test this hypothesis, we leveraged a well-established method to monitor glucose efflux from the lysosome by loading the lysosome with sucrose followed by the delivery of invertase via endocytosis to hydrolyze sucrose into glucose and fructose (Figure 6C)^45^. Coupling this method with LysoIP enabled us to assess whether glucose is trapped in the lysosome (Figure 6C and 6D). To this end, we adapted and validated a derivatization method that increases the sensitivity of hexose detection in purified lysosomes using LC-MS analysis^46^ (Figures 6D, S6C and S6D). Consistent with SLC45A1 functioning as a sugar transporter from the lysosome, we observed a significant accumulation of glucose in lysosomes derived from *SLC45A1*-KO cells compared to those from wildtype or rescued cells when treated with sucrose and invertase (Figures 6E and F). To validate these findings *in vivo*, we generated wildtype and *Slc45a1*-KO mice expressing LysoTag under the *Syn1* promoter, enabling robust purification of neuronal lysosomes (Figures 5G and S6E). Strikingly, we found a significant accumulation of hexose sugars in lysosomes derived from *Slc45a1*-/- neurons compared to those from wildtype mice (Figures 6H and 6I). These results establish a critical role for SLC45A1 in the clearance of hexose sugars from neuronal lysosomes, shedding light on its essential function in maintaining lysosomal homeostasis and energy metabolism in neurons.

**Figure 6.**
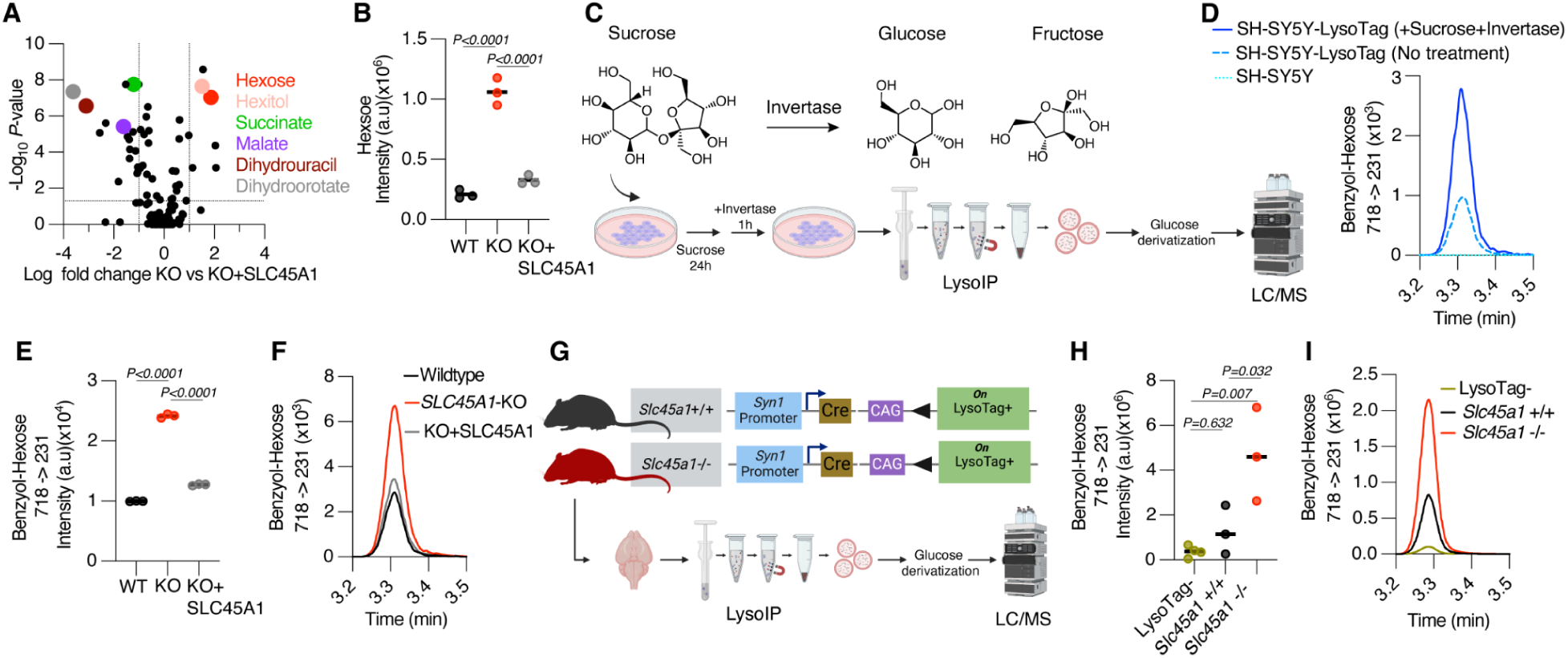
SLC45A1 deficiency leads to sugar accumulation in the lysosome. (A) Untargeted metabolomic analysis of *SLC45A1*-KO and rescued (KO+SLC45A1) SH-SY5Y cells under 72 h serum deprivation conditions. Data is presented as a volcano plot of log_2_-transformed fold changes in the abundance of putatively identified metabolites between *SLC45A1*-KO and rescued cells. The horizontal line indicates a *P*-value of 0.05 and the vertical lines indicate a fold change of 2. *n* = 3 biological independent samples per genotype. *P*-values were calculated by one-way ANOVA with Tukey’s HSD test. Metabolomics data for wildtype, *SLC45A1*-KO, and rescued cells are provided in Table S10. (B) Targeted quantitation of hexose (C_6_H_12_O_6_) in wildtype, *SLC45A1*-KO, and rescued cells from A. *P*-values were calculated using one-way ANOVA with Tukey’s HSD test. Horizontal line represents the mean (*n* = 3 biologically independent samples). (C) Schematic overview of the procedure for loading lysosomes with sucrose to monitor glucose efflux. Cells were treated with sucrose (50 mM) for 24 h, followed by treatment with invertase (0.5 mg/ml) for 1 h. Hexose in purified lysosomes was derivatized to benzoyl-hexose for measurement by LC-MS/MS analysis. (D) Representative extracted ion chromatogram of benzoyl-hexose measurement in purified lysosomes following sucrose and invertase treatment compared to untreated condition, indicating hexose enrichment in lysosomes after treatment. (E) Measurement of benzoyl-hexose from purified lysosomes in wildtype, *SLC45A1*-KO, and rescued SH-SY5Y cells upon treatment with sucrose and invertase. Horizontal line represents the mean (*n* = 3 biologically independent samples). *P*-values were calculated using one-way ANOVA with Tukey’s HSD test. (F) Representative extracted ion chromatogram of wildtype, *SLC45A1*-KO, and rescued cells from the same experiment in E. (G) Schematic overview for analyzing lysosomal hexose specifically from neurons using LysoIP. *Syn1* promoter-driven Cre recombinase allows LysoTag expression in neurons of *Slc45a1*+/+ or *Slc45a1*-/- mice. Control mockIP was done using Cre-negative mice (LysoTag-). (H) Measurement of derivatized hexose from purified lysosomes of mouse brain neurons. LysoIP was performed in *Slc45a1*+/+;*Syn1*-Cre+;LysoTag+ (*n* = 3), *Slc45a1*-/-;*Syn1*-Cre+;LysoTag+ (*n* = 3) and LysoTag-controls for mockIP (*n* = 4). Horizontal line represents the mean. *P*-values were calculated using one-way ANOVA with Tukey’s HSD test. (I) Representative extracted ion chromatogram from each LysoIP in H.

**Figure S6.**
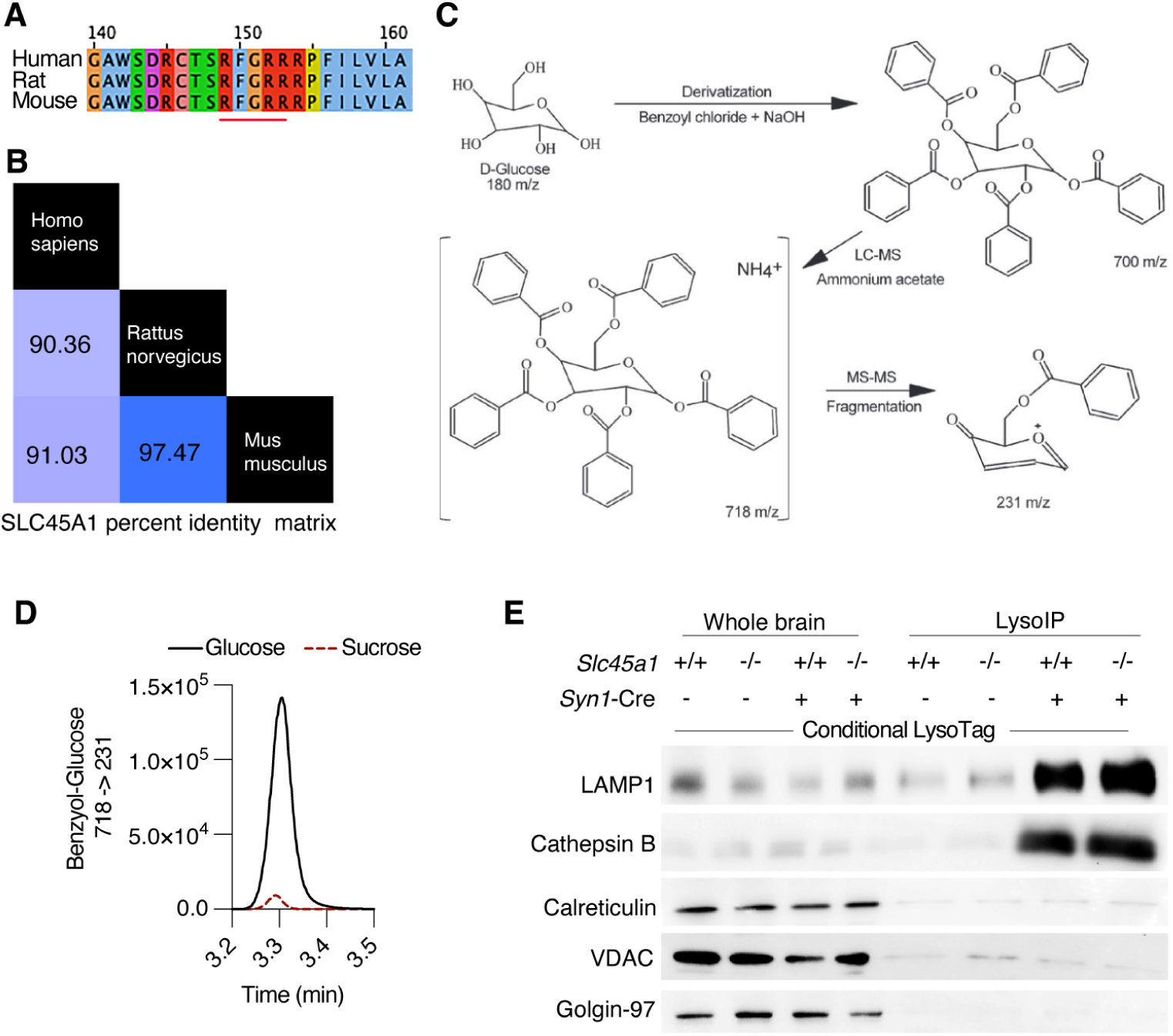
Characterization of SLC45A1 protein as lysosomal hexose transporter. (A) Protein sequence alignment of human, rat and mouse SLC45A1 by Jalview version 2.11.3.3. The red line marks the conserved RXGRR sequence motif found in sugar transporters. (B) The percent identity matrix, generated with Clustal2.1, demonstrates the high conservation of the SLC45A1 protein among human, rat, and mouse. (C) Overview of hexose derivatization to increase detection sensitivity. Hexose is derivatized to form benzoyl-hexose. Samples were analyzed for the parent ion at 718 *m/z* [M^+^NH_4_]^+^ and the product ion at 231 *m/z* using LC-MS/MS. (D) Extracted ion chromatogram indicating that after derivatization of 100 µM glucose and sucrose only benzoyl-glucose is detected as a product at 231 m/z. (E) LysoIP was used to isolate neuronal lysosomes from the brains of *Slc45a1*+/+ and *Slc45a1***-/-** mice. Tmem192-3xHA is LysoTag. LAMP1 and Cathepsin B were used as lysosomal markers, while Golgin-97, VDAC1, and Calreticulin were used as markers for Golgi, mitochondria, and endoplasmic reticulum, respectively.

## Discussion

Lysosomal dysfunction is implicated in a wide range of neurodegenerative diseases and many of the lysosomal storage disorders (LSDs) manifest with severe neurological symptoms^5,10,12^. Despite their critical role in the brain, how lysosomal composition and function differ across various brain cell types remains poorly understood.

Using the LysoIP method coupled with high-resolution mass spectrometry^13^, we created the first comprehensive lysosomal protein atlas across major brain cell types, including neurons, astrocytes, microglia, and oligodendrocytes. This was achieved by leveraging the LysoTag mouse model, bred with mice expressing Cre recombinases under cell-type-specific promoters^13^. Our brain lysosomal atlas revealed both well-characterized and novel lysosome-associated proteins, expanding our understanding of lysosomal function and diversity in the brain. Although lysosomal composition was largely conserved, we observed distinct cell-type-specific lysosomal signatures, suggesting specialized lysosomal functions in different brain cell types. Notably, several proteins associated with LSDs displayed differential expression patterns across cell types, suggesting distinct susceptibilities to lysosomal dysfunction in different brain cell types and regions^47^. While lysosomal specialization, involving both membrane-associated proteins and lumenal enzymes, appears to be primarily driven by transcriptional programs specific to each cell type, our data also highlight how cell-type-resolved lysosomal profiling in a complex organ like the brain can uncover additional insights related to protein trafficking between cell types^48^. Further characterization of these protein exchange events under both homeostatic and disease conditions could reveal novel mechanisms regulating protein turnover and intercellular communication in the brain through lysosomal pathways.

The identification of SLC45A1 as a neuron-specific lysosomal protein highlights the effectiveness of LysoIP in uncovering cell-type-specific lysosomal components. To our knowledge, SLC45A1 is the first lysosomal protein identified as specific to neurons. Mutations in SLC45A1 cause a monogenic neurological disease characterized by epilepsy and intellectual disability^14–16^. Here, we show that SLC45A1-associated disease represents a previously uncharacterized lysosomal storage disorder. Our data demonstrate that loss of SLC45A1 leads to hexose accumulation in lysosomes, consistent with a sugar transport function, along with increased LAMP1 levels, expansion of lysosomal compartment and disruption of its ultrastructure, altered autophagy, and reduced proteolytic activity—hallmarks of LSDs. While our results confirm SLC45A1’s role in sugar transport from the lysosome and not on the plasma membrane as previously thought^14,34^, it will be important to further define its endogenous substrates, which may include sugar moieties from degraded glycoproteins and glycolipids^3^. Additionally, it remains to be understood why neurons have a specific lysosomal sugar transporter and whether similar transport mechanisms exist in other brain cell types.

Beyond its transport function, SLC45A1 interacts with V1 subunits of the V-ATPase complex, which is crucial for maintaining lysosomal acidity^49^. Lysosomal pH regulation is essential for proper lysosomal function, including catabolism, metabolite export and signalling^49^, and this regulation is disrupted in SLC45A1-deficient lysosomes, implicating SLC45A1 in stabilizing V-ATPase. Indeed, loss of SLC45A1 leads to a significant reduction of the V1 subunits on the lysosomal membrane. Given that nutrients including glucose are known to regulate V-ATPase activity and assembly^38,39^, we speculate that SLC45A1’s role in sugar transport may influence V-ATPase activity and, by extension, lysosomal pH and metabolic signaling^50^. Understanding the intersection between SLC45A1-mediated sugar transport and V-ATPase regulation could shed light on broader mechanisms of the lysosomal function in the neurons.

Mitochondrial dysfunction is a well-established pathological mechanism in LSDs^5^. Recent studies suggest that disruption of lysosomal acidity can lead to iron deficiency, as iron is trafficked through the lysosome, which in turn disrupts mitochondrial complexes containing iron-sulfur clusters^41,42^. In line with these findings, we observed that SLC45A1 deficiency impacts cellular iron homeostasis, elevating reactive oxygen species (ROS) and leading to mitochondrial dysfunction. This suggests that SLC45A1-associated lysosomal dysfunction is linked to broader metabolic disruptions in neurons. Given the high metabolic demand of neurons, which rely heavily on mitochondrial activity for survival, these findings warrant further investigation into the interplay between lysosomal function and energy balance in neuronal cells^51^.

In conclusion, our lysosomal atlas provides a valuable resource for understanding the diverse roles of lysosomes across brain cell types and how cell-type-specific lysosomal functions contribute to brain diseases. The identification of SLC45A1 as a key regulator of lysosomal pH and sugar export enhances our understanding of the complexity and specificity of lysosomal function, particularly in neurons. These findings open new avenues for studying lysosomal biology in the brain and its implications for neurodegenerative diseases and lysosomal storage disorders.

## Acknowledgments

We thank all members of the Abu-Remaileh and Ori laboratories. We would like to thank the Zuchero and Gibson labs at Stanford University for providing primary oligodendrocyte and primary microglia, respectively. The authors gratefully acknowledge support from the FLI Core Facilities Proteomics, and Imaging and would like to thank Ivonne Heinze and Maleen Hofmann for their excellent technical assistance. We also thank the Metabolomics Knowledge Center at Stanford Sarafan ChEM-H Institute (directed by Dr. Yuqin Dai) and the Transgenic Knockout and Tumor Model Center at Stanford University. This work was supported by grants from Beatbatten, the NCL Foundation (NCL-Stiftung), the NIH Director’s New Innovator Award Program (1DP2CA271386), the Knight Initiative for Brain Resilience at Stanford University, the Chan Zuckerberg Initiative Neurodegeneration Challenge Network (2024-338545), and Aligning Science Across Parkinson’s [ASAO-000463] through Michael J. Fox Foundation for Parkinson’s Research (MJFF) to M.A-R. A.O. is supported by the German Research Council (Deutsche Forschungsgemeinschaft, DFG) via the Research Training Group ProMoAge (GRK 2155), and the Chan Zuckerberg Initiative Neurodegeneration Challenge Network (award numbers: 2020-221617, 2021-230967 and 2022-250618). A.O. and J.C.H. are supported by the Fritz-Thyssen Foundation (award number: 10.20.1.022MN), and the NCL Stiftung. The FLI is a member of the Leibniz Association and is financially supported by the Federal Government of Germany and the State of Thuringia. K.N. is supported by the Sarafan ChEM-H Chemistry/Biology Interface Program as Kolluri Fellow. K.N is additionally supported by the Bio-X Stanford Interdisciplinary Graduate Fellowship affiliated with the Wu Tsai Neurosciences Institute (Bio-X SIGF: Mark and Mary Steven’s Interdisciplinary Graduate Fellow). M.A-R is a Stanford Terman Fellow and a Pew-Stewart Scholar for Cancer Research, supported by the Pew Charitable Trusts and the Alexander and Margaret Stewart Trust.

## Author contributions

Conceptualization: MA-R, AO, AG, JCH

Data curation: AO, JCH, AG

Investigation: AG, JCH

Methodology: MA-R, AO, AG, JCH, UNM, NNL, MG, KN, NGO

Project administration: MA-R, AO

Data analysis: JCH, AG, DDF, ESR, UNM, WD, KN, AO

Supervision: MA-R, AO

Visualization: JCH, AG, AO

Writing – original draft: MA-R, AO, JCH, AG

Writing – review & editing: All authors

Both AG and JCH contributed equally and have the right to list their name first in their CV.

## Declaration of interest

M.A-R. is a scientific advisory board member of Lycia Therapeutics and advisor for Scenic Biotech. All other authors declare no competing interests.

## List of Supplemental Tables provided as separate Excel tables

Table S1: Differential protein abundance against mockIP and lists of protein definitions

Table S1A: Known lysosomal proteins; related to Figures 2A, 2C, 3C.
Table S1B: Core lysosomal proteins; related to Figures S1C and S2B.
Table S1C: Nuclear proteins; related to Figure S2B.
Table S1D: Differential protein abundance against mockIP for different Cre lines; related to Figure 1C.

Table S2: Definition of LysoIP-enriched proteins across Cre lines

Table S2A: Classifier Total Brain CMV-Cre; related to Figure 2A.
Table S2B: Classifier Neurons Syn1-Cre; related to Figure 2A.
Table S2C: Classifier Astrocytes Gfap-Cre; related to Figure 2A.
Table S2D: Classifier Oligodendrocytes Olig2-Cre; related to Figure 2A.
Table S2E: Classifier Microglia Cx3cr1-Cre; related to Figure 2A.
Table S2F: Ranked LysoIP-enriched proteins across all classifiers; related to Figure 2C and S2D.

Table S3: Enrichment of GO terms in LysoIP-enriched proteins

Table S3A: GO Cellular Compartments; related to Figure 2B.
Table S3B: GO Biological Processes; related to Figure S2C.
Table S3C: GO Molecular functional; related to Figure S2C.

Table S4: Differential abundance of LysoIP-enriched proteins across Cre lines; related to Figure 2A, 2C, 3B, 3C, 3D, 3F, 3G, S3B, S3C, S3D.

Table S5: Comparison between scRNAseq and LysoIP proteomics; related to Figure 3E, S3E, S3F, S3G and S3H.

Table S6: Reference peptides for SLC45A1 targeted proteomics; related to Figure S5B and S5F.

Table S7: Untargeted lipidomics of wildtype, *SLC45A1*-KO, KO+SLC45A1 (rescued cells) SH-SY5Y cells under serum starvation; related to Figure 5D.

Table S8: BioID for SLC45A1

Table S8A: BioID candidates table; related to Figure 5I.
Table S8B: BioID report table; related to Figure 5I.

Table S9: Quantification of V-ATPase subunits in LysoIPs from SH-SY5Y cells; related to Figure 5K.

Table S9A: V-ATPse quantification in LysoIPs related to Figure 5K.
Table S9B: V-ATPse quantification in corresponding whole cell lysates; related to Figure 5K.

Table S10: Untargeted metabolomics of wildtype, SLC45A1-KO, and KO+SLC45A1 (rescued cells) SH-SY5Y cells under serum starvation; related to Figure S5N, 6A and 6B.

## Materials and Methods

Antibodies, plasmids, chemicals, oligonucleotides, and software are listed in the key resources table.

### Key resources table

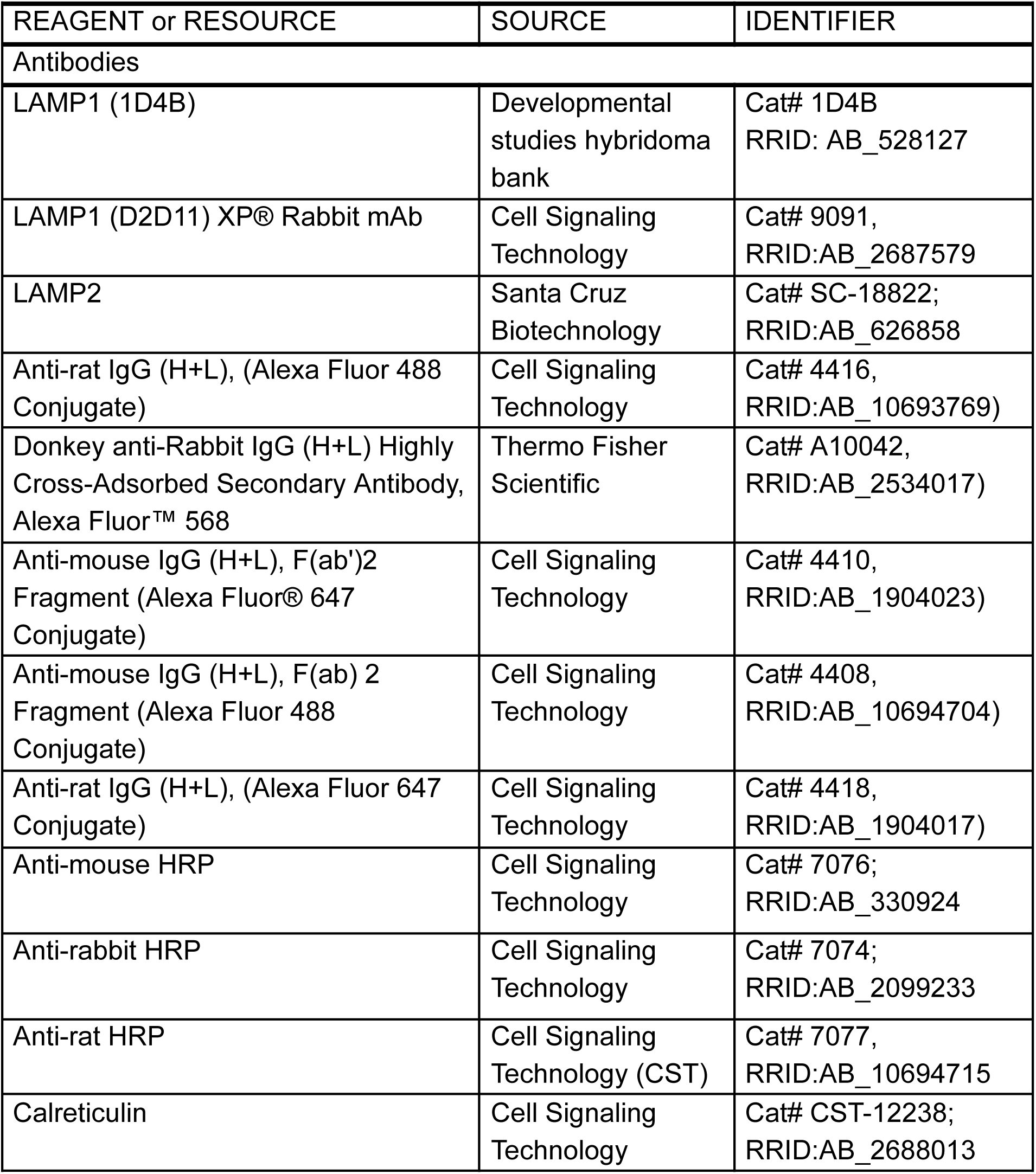
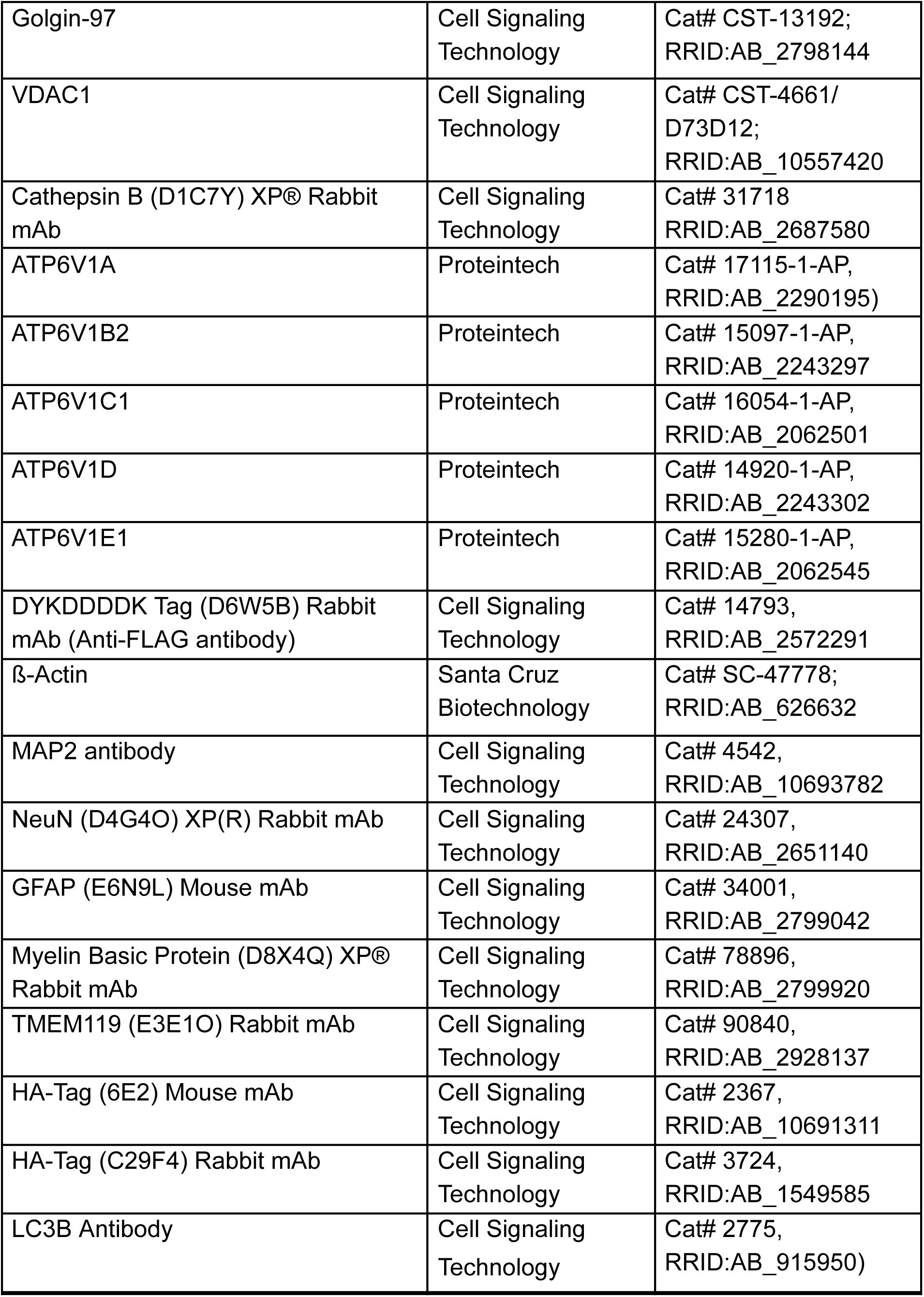
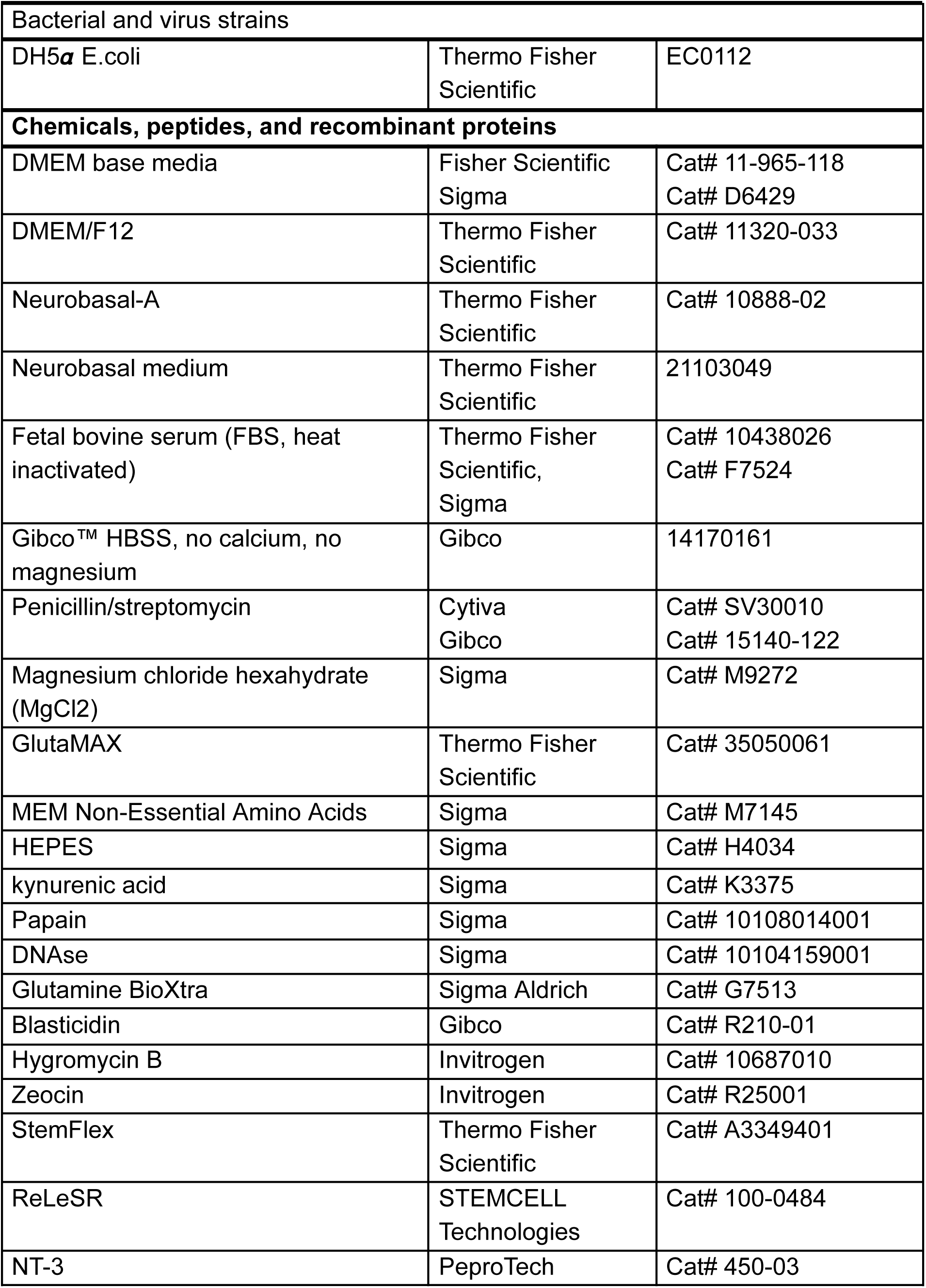
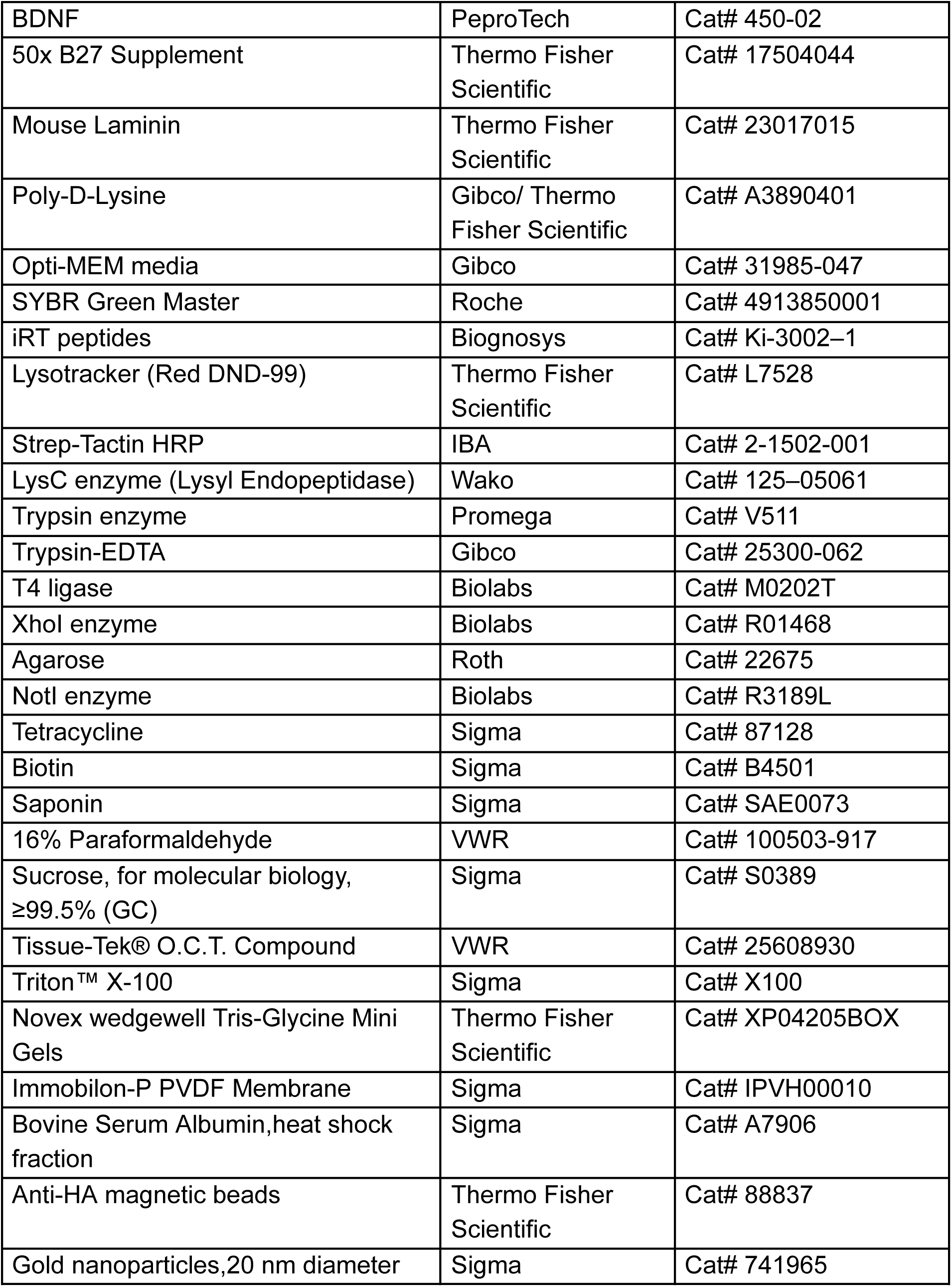
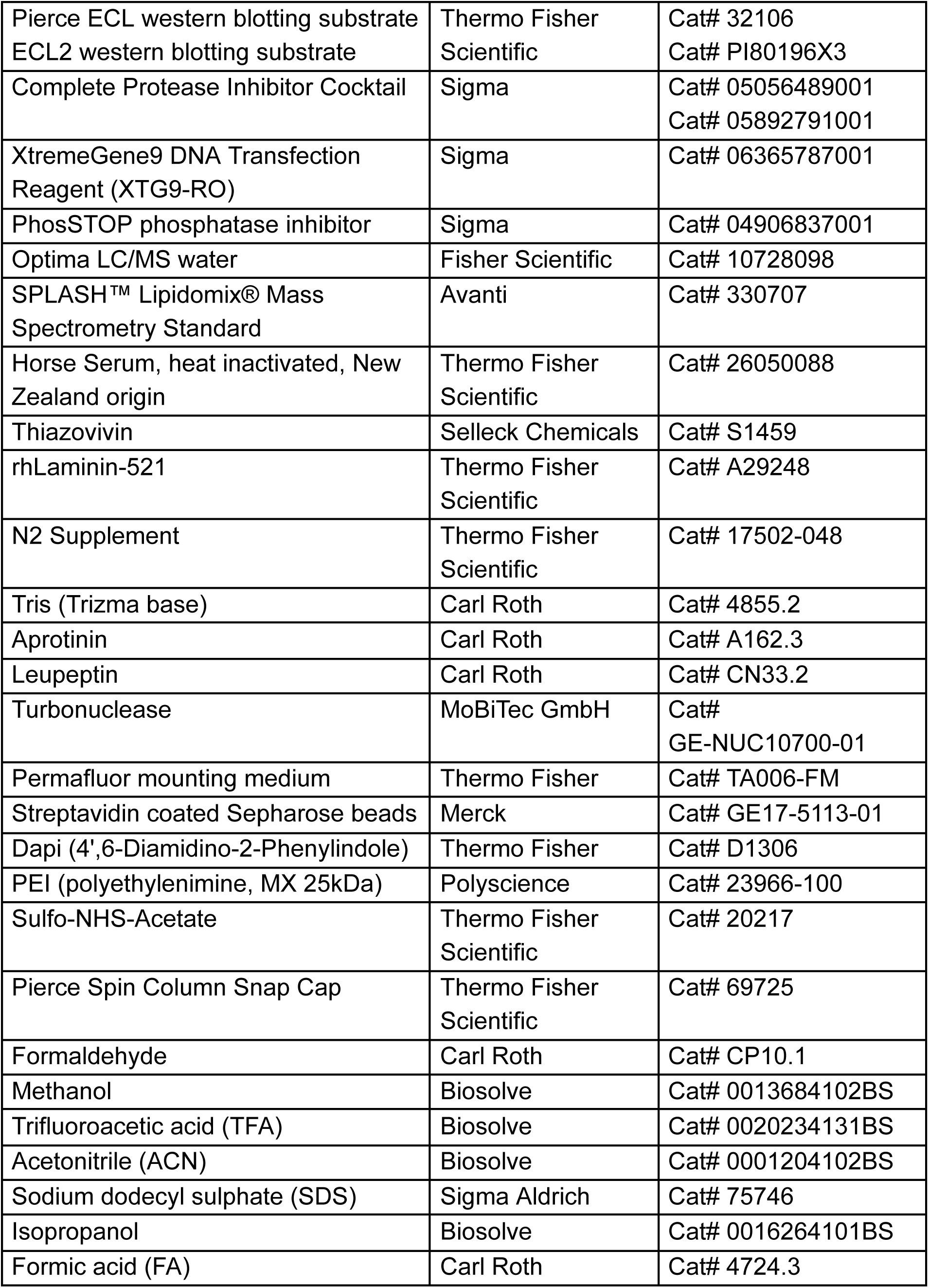
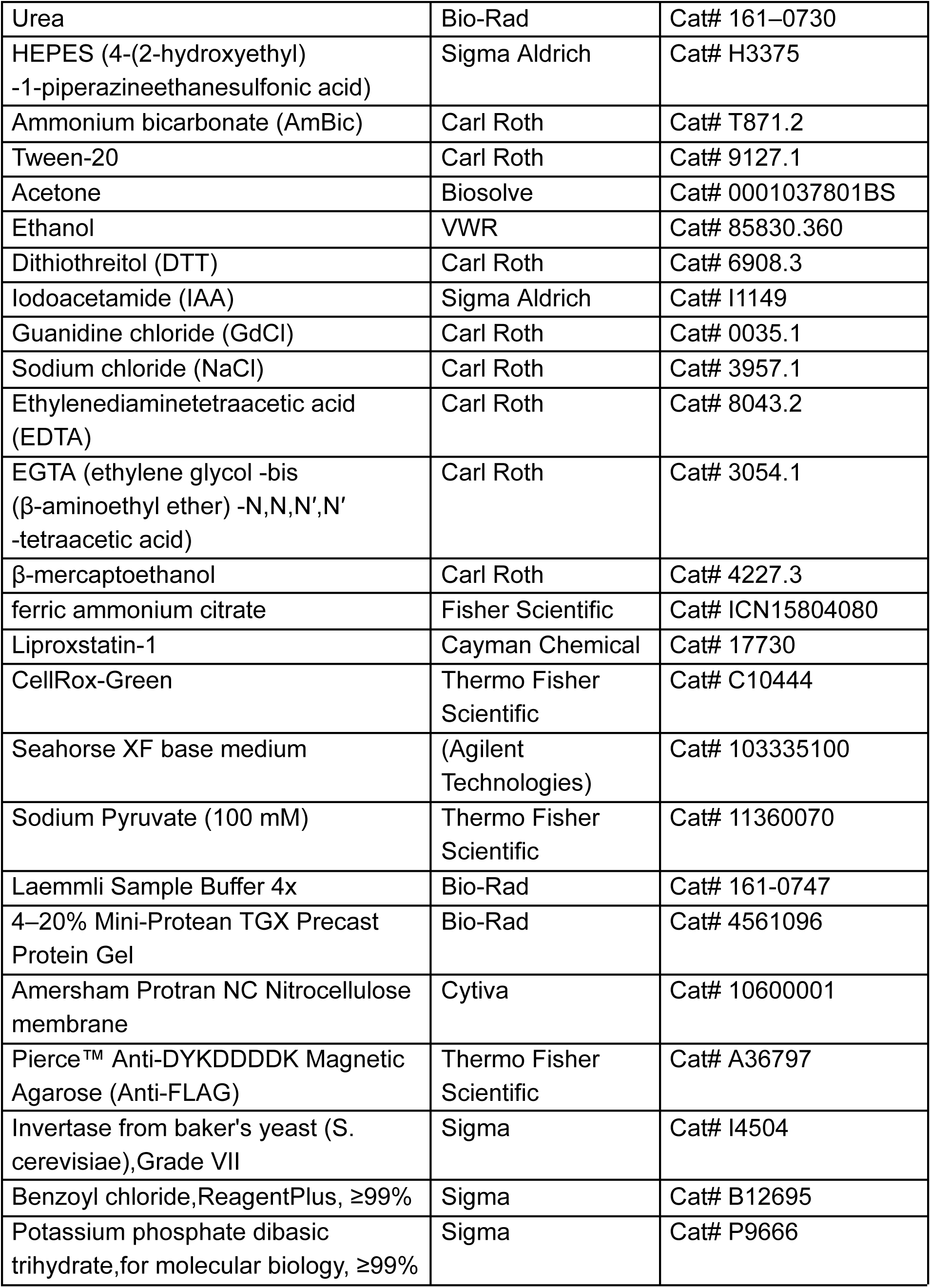
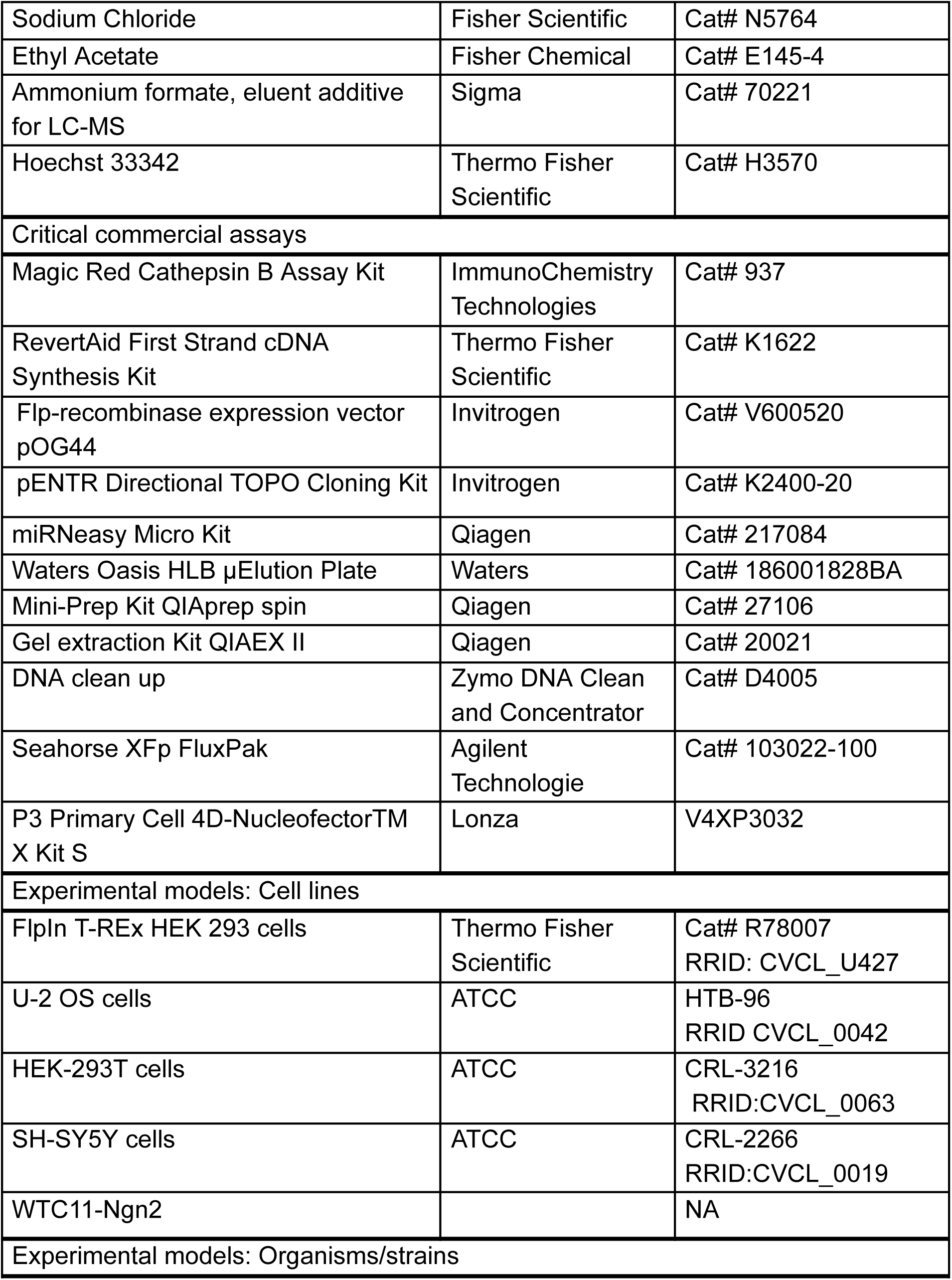
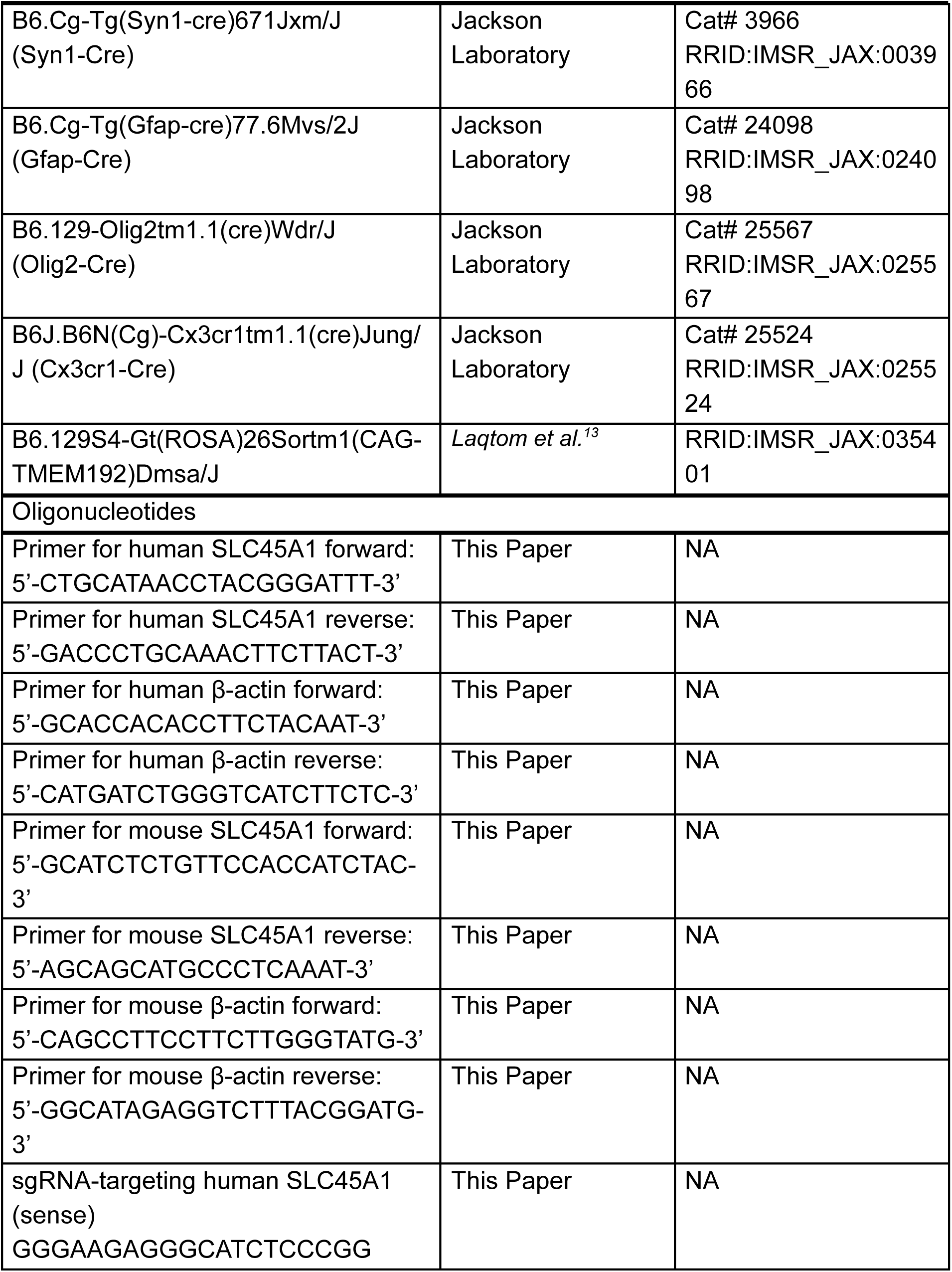
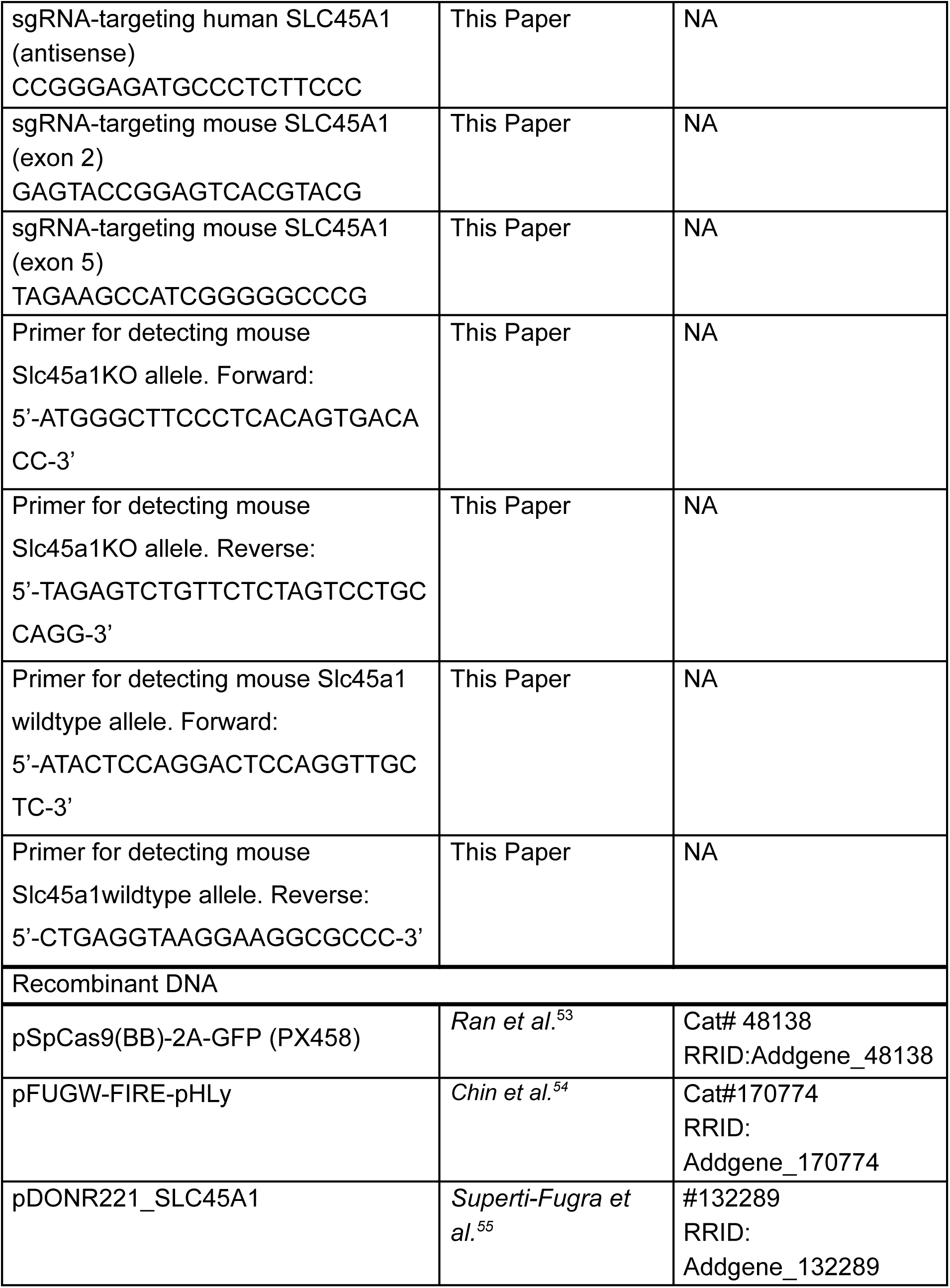
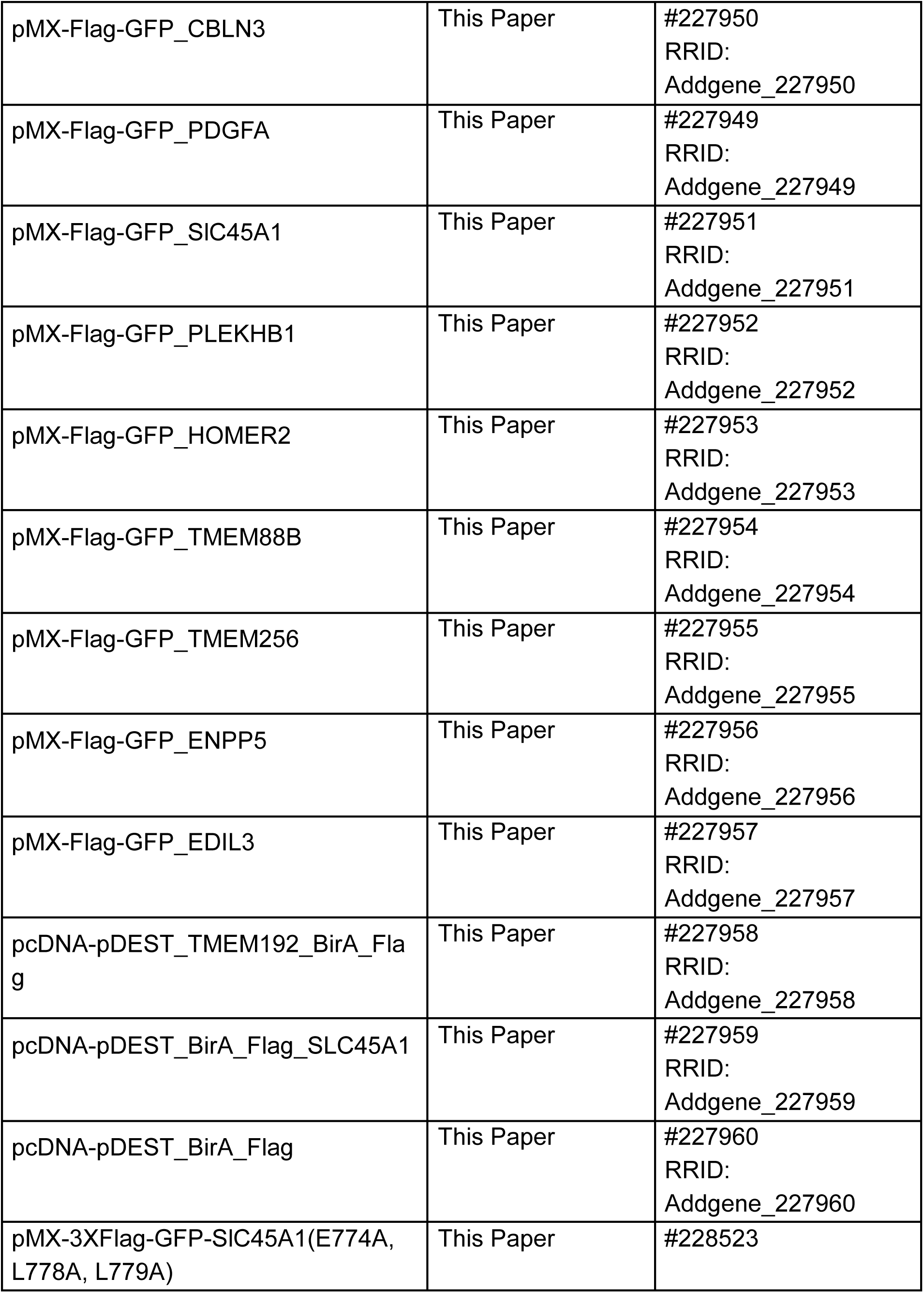
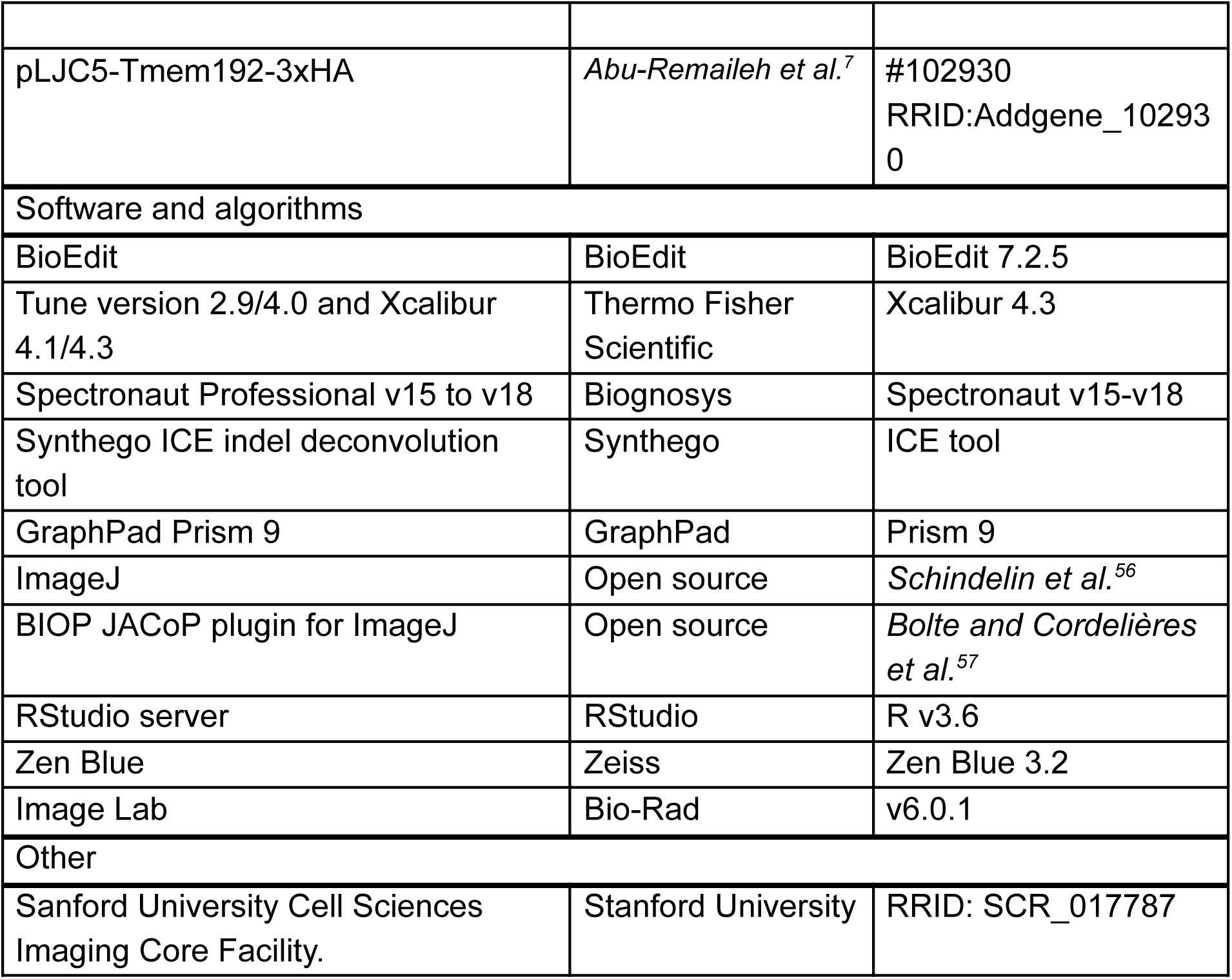

### Mouse studies

Mice were maintained on a standard light–dark cycle with ad libitum access to food and water in a room with controlled temperature (22°C) and humidity (around 50%). Cages were cleaned every 4–5 days, and water and food supplies were checked daily.

To induce LysoTag expression specifically in each of the four major brain cell types, *Rosa26;lox-stop-lox-TMEM192-3×HA* mice^13^ were crossed with mice expressing Cre recombinase driven by specific promoters: neuronal-specific promoter *Syn1* (B6.Cg-Tg(*Syn1*-Cre)671Jxm/J, Stock No: 3966), astrocyte-specific promoter Gfap (B6.Cg-Tg(*Gfap*-Cre)77.6Mvs/2J, Stock No: 24098), oligodendrocyte-specific promoter Olig2 (B6.129-*Olig2tm1*.1(Cre)Wdr/J, Stock No: 25567) and microglia-specific promoter Cx3cr1 (B6J.B6N(Cg)-*Cx3cr1tm1.1*(Cre)Jung/J, Stock No: 25524). All Cre-driver mice were purchased from The Jackson Laboratory.

*Slc45a1*-/- mice were generated by the Transgenic Knockout and Tumor Model Center at Stanford University using CRISPR/Cas9 technology, targeting a 5.8 kb (5837 bp) region for deletion between exon 2 and exon 5 in the *Slc45a1* gene (NM_173774.3, chr4, 150713853-150736631). The guide RNAs used were 5’-GAGTACCGGAGTCACGTACG-3’ (exon 2) and 5’-TAGAAGCCATCGGGGGCCCG-3’ (exon 5). To detect KO allele in mice, the primers used are 5’-ATGGGCTTCCCTCACAGTGACACC-3’ (forward) and 5’-TAGAGTCTGTTCTCTAGTCCTGCCAGG-3’ (reverse), with an expected PCR product size of 664 bp. For the WT allele, the primers are 5’-ATACTCCAGGACTCCAGGTTGCTC-3’ (forward) and 5’-CTGAGGTAAGGAAGGCGCCC-3’ (reverse), with an expected product size of 400 bp. To generate neuron-specific LysoTag expression in *Slc45a1*+/+ or *Slc45a1*-/- mice, we crossed *Slc45a1*+/-; *Syn1*-Cre+/- mice with *Slc45a1*+/-;LysoTag (homozygous) mice. All the procedures involving mice were carried out in accordance with the approved guidelines by the Institutional Animal Care and Stanford University (APLAC-33464).

### Cell culture

HEK-293T, and SH-S5Y5 (a kind gift from Gitler lab, Department of Genetics, Stanford University School of Medicine) cells were cultured in DMEM base media (Fisher Scientific) supplemented with 2 mM glutamine, 10% heat inactivated fetal bovine serum (FBS, Fisher Scientific) and 1% penicillin/streptomycin (Cytiva). For serum depletion, SH-S5Y5 cells were cultured in 24, 48, and 72 h in DMEM base media supplemented with 2 mM glutamine, 1% penicillin/streptomycin. U2OS cells (a kind gift from Pospiech lab, Leibniz Institute on Aging-Fritz Lipmann Institute) cells were cultured in DMEM base media (Sigma) supplemented with 2 mM glutamine (Sigma), 10% heat inactivated FBS (Sigma) and 1% penicillin/streptomycin (Gibco). Flp-In-T-REx HEK 293 cells, referred to as Flp-HEK293, expressing BirA*-SLC45A1, TMEM192-BirA* or BirA* (negative control) were generated as described in the material and methods section “Proximity labeling and sample preparation for BioID”. Cells were grown in DMEM (Sigma) supplemented with 10% (v/v) heat inactivated FBS (Sigma), 2 mM L-Glutamine (Sigma), 15 µg/ml Blasticidin and 100 µg/ml Hygromycin B. The parental Flp-HEK293 cell line was grown in presence of 100 µg/ml Zeocin and 15 µg/ml Blasticidin. Upon generation of stable cell lines, Zeocin was replaced by 100 µg/ml Hygromycin B. All the cells were maintained at 37°C, 5% CO_2_ and 95% humidity in a CO_2_ incubator.

### Human iPSCs and iPSC-derived neurons

In this study, male WTC11 human induced pluripotent stem cells (iPSCs)^58^ engineered to express NGN2 under a doxycycline-inducible system at the AAVS1 safe harbor locus^52,59^, were used following the ethical guidelines of the Stem Cell Research Oversight (SCRO) (SCRO-845). The iPSCs were maintained in StemFlex™ Medium (Thermo Fisher Scientific) on plates coated with rhLaminin (Thermo Fisher Scientific), diluted in 1x DPBS to a final concentration of 2.5 μg/ml. To initiate differentiation^59^ the iPSCs were detached using ReLeSR™ (StemCell Technologies), and the resulting cell pellet was resuspended in N2 Pre-Differentiation Medium containing Knockout DMEM/F12 (Thermo Fisher Scientific), 1x MEM Non-Essential Amino Acids (Sigma), 1x N2 Supplement (Thermo Fisher Scientific), 10 ng/ml NT-3 (PeproTech), 10 ng/ml BDNF (PeproTech), 1 μg/ml Mouse Laminin (Thermo Fisher Scientific), 10 nM Thiazovivin (ROCK inhibitor) (Selleck Chemicals) and 2 μg/ml doxycycline hydrochloride to induce NGN2 expression. The iPSCs were seeded into rhLaminin-coated 6-well plates at a density of 1.5×10⁶ cells per well and cultured in 2 ml of N2 Pre-Differentiation Medium. After three days (Day 0), pre-differentiated cells were dissociated with Accutase, centrifuged, and resuspended in Classic Neuronal Medium. This medium consisted of a 1:1 mixture of DMEM/F12 (Thermo Fisher Scientific) and Neurobasal-A (Thermo Fisher Scientific) as the base, supplemented with 1x MEM Non-Essential Amino Acids, 0.5x GlutaMAX (Thermo Fisher Scientific), 0.5x N2 Supplement, 0.5x B27 Supplement (Thermo Fisher Scientific), 10 ng/ml NT-3, 10 ng/ml BDNF, 1 μg/ml Mouse Laminin, and 2 μg/ml doxycycline hydrochloride. The cells were counted and seeded at a density of 3.0×10⁵ cells per well in a Poly-D-Lysine coated 12-well plate with 2 ml of Classic Neuronal Medium, or 1.5×10⁵ cells per well in a Poly-D-Lysine coated 24-well plate with 1 ml of medium. On Day 7, half of the medium was replaced with fresh Classic Neuronal Medium without doxycycline, and the cells were cultured for an additional 7 days. On Day 14, another half-medium change was performed with the same medium, and the cells were further cultured for another 7 days. On Day 21, the differentiated cells were harvested for subsequent experiments. The detailed protocol can be found at^60^.

### Endogenous N-terminal 3x FLAG Tagging of SLC45A1 in iPSCs

Endogenous N-terminal 3x FLAG tagging was performed on the WTC11-Ngn2 iPSC line, with the FLAG tag inserted immediately after the start codon. Twenty-four hours before nucleofection, iPSCs (WTC11-Ngn2) were pretreated with 10 µM thiazovivin (ROCK1 inhibitor). On the day of transfection, the ribonucleoprotein (RNP) complex was prepared by combining 6 µg (0.036 nmol) of Cas9 enzyme (IDT Alt-R™ S.p. Cas9 Nuclease V3) with 3.2 µg (0.15 nmol) of sgRNA (Synthego - 2’-O-methyl modified). The mixture was incubated for 15 min at 37°C. iPSCs were dissociated using ReLeSR™. 0.3×10^6^ live cells were resuspended with the RNP complex and 20 µL of P3 Primary Cell Nucleofector Solution (Lonza), following the manufacturer’s instructions. The resuspended cells were transferred to one well of a 16-well Nucleocuvette Strip. The donor template for 3x FLAG consisted of a 200-bp-long ssODN: agcagcccagcggggacagggatgcctgccgtctccacccacagggacgcccaccagccctccccacgATGGACTACAAAGACCATGACGGTGATTATAAAGATCATGACATCGATTACAAGGATGACGATGACAAGATCCCCGCAGCCAGCAGCACCCCGCCGGGAGATGCCCTCTTCCCCAGCGTGGCCCCACAGGAC (IDT Alt-R HDR donor oligonucleotide). The template included a left homology arm (71 bp), the 3x FLAG tag sequence: GACTACAAAGACCATGACGGTGATTATAAAGATCATGACATCGATTACAAGGATGACGATGACAAG (66 bp), and a right homology arm (63 bp). Lowercase letters indicate non-coding sequence. The template was designed so that only 10 bp of the 19-bp guide sequence: CTGGCTGCGGGGATCATcg were present, preventing cleavage of the template by the RNP complex before or after integration. 6 µg (0.1 nmol) of ssODN were delivered into the cells alongside the RNP complex. Nucleofection was performed in a 4D-Nucleofector (Lonza) using the CA137 program. Nucleofected iPSCs were plated in a 24-well plate in the presence of AZD7648 at a final concentration of 1 µM, along with continued thiazovivin treatment. After 24 h of incubation, the cells were switched to fresh growth medium without AZD7648 but continued thiazovivin treatment^61^.

### Plasmid construction

Gene fragments for lysosomal localization validation where ordered from TWIST Bioscience (Gene fragments without Adapters, codon optimized; encoding proteins: EDIL3, ENPP5, PDGFA, PLEKHB1), and IDT (gblocks; encoding proteins: TMEM88B, TMEM256, CBLN3, HOMER2). For SLC45A1, pDONR221_SLC45A1 was a gift from RESOLUTE Consortium & Giulio Superti-Furga (Addgene plasmid #132289; http://n2t.net/addgene:132289; RRID:Addgene_132289) used as template. Candidate proteins were cloned into a pMXs-IRES-Blast-3xFlag-GFP vector by restriction enzyme cloning strategy (XhoI and NotI). After DNA clean up (Zymo DNA Clean and Concentrator, following manufacturer’s instruction), the gene fragments were ligated using T4 ligase (Biolabs) and transformed into DH5α E.coli (Thermo Fisher)^62^. DNA extraction (QIAprep spin) was followed by sequencing (Microsynth) and sequence validation (BioEdit) before further use of the plasmid. All plasmids are available at Addgene.

### Virus production and transduction

HEK293T cells were transfected with retroviral plasmids along with packaging plasmids and VSV-G envelope using XtremeGene9 transfection reagent (Sigma). After 16 h, the culture medium was replaced with DMEM supplemented with 30% inactivated fetal bovine serum. After 48 h, the supernatant was harvested, centrifuged for 5 min at 300 g to remove cells, and frozen at -80°C. Stable expression cell lines were prepared by first plating 1×10^6^ SH-SY5Y cells were seeded in 6-well plates in DMEM with 10% inactivated fetal bovine serum, 8 μg/ml polybrene, and 100-250 µl of virus-containing media. Spin infection was then performed at 2,200 rpm for 45 min at 37°C. Following a 16 h incubation, virus-containing medium was replaced with fresh culture medium.

### Generation of knockout cell lines using CRISPR–Cas9 technology

The following SLC45A1-targeting sgRNA: sense, 5’-caccgGGGAAGAGGGCATCTCCCGG-3’ and antisense, 5’-aaacCCGGGAGATGCCCTCTTCCCc-3’ oligonucleotides were cloned into the pX458 vector. pX458-sgRNA plasmids were then transfected into SH-SY5Y cells using the XtremeGene 9 (1:3) transfection reagent. 48 h post-transfection, GFP-positive (encoded on pX458) cells were single-sorted into 96-well plates, each well containing 200 μl of DMEM with 30% FBS and penicillin/streptomycin. The single-sorted cells were incubated at 37°C and 5% CO_2_ to form colonies. Once the colonies formed, they were harvested, and the sgRNA target sites were PCR-amplified and sequenced using Sanger sequencing. Indels at the sgRNA target sites were analyzed using the Synthego ICE indel deconvolution tool (https://synthego.com).

### Rapid isolation of lysosomes (LysoIP) from brain tissue and cells

Whole brains were collected on an ice-cold plastic dish from mice after euthanasia (2-3 months old mice, mixed gender). Fresh brain tissue was used immediately without freezing, following steps similar to the original LysoIP protocol^7,13^. Each mouse was processed separately to ensure rapid isolation of lysosomes using pre-chilled equipment and buffers prepared in Optima LC/MS water. Each brain was gently homogenized in 950 µl phosphate-buffered saline (PBS, pH ∼7.4) containing Complete EDTA-free Protease Inhibitor Cocktail (Roche). The homogenization was done with 25 strokes in 2 ml homogenizer. The liquid fraction was then transferred to a 2 ml tube, and 25 µl (equivalent to 2.5% of the total tissue) was reserved for further processing of the whole brain fraction. To minimize contamination from the pellet in the next step, an additional 1 ml of PBS with Protease Inhibitor Cocktail was added to the tube. The homogenate was centrifuged at 1000 g for 2 min at 4°C. The supernatant, containing cellular organelles including lysosomes, was incubated with 100 µl of pre-washed anti-HA magnetic beads (Thermo Fisher Scientific) in PBS (with Protease Inhibitor Cocktail) on a gentle rotator shaker for 15 min at 4°C. Immunoprecipitates were washed three times with 1 ml PBS (with Protease Inhibitor Cocktail) on a DynaMag Spin Magnet. For protein extraction from lysosomes, the beads with bound lysosomes were resuspended in 80 µl ice-chilled 1x extraction buffer (50 mM HEPES at pH 7.4, 40 mM NaCl, and 2 mM EDTA, 1% Triton X-100 (Sigma), 1.5 mM sodium orthovanadate (NaVO_4_), 30 mM sodium fluoride (NaF), 10 mM sodium pyrophosphate (Na_4_P_2_O_7_), 10 mM sodium β-glycerophosphate and Complete EDTA-free Protease Inhibitor Cocktail). The mixture was incubated for 15 min at 4°C, after which the beads were removed using a magnet. For the whole brain sample, 160 µl ice-chilled 1x extraction buffer was added, followed by brief vortexing to extract cellular proteins. After a 15 min incubation, the protein extract was centrifuged at maximum speed for 15 min at 4°C, and the supernatant was collected. For SH-SY5Y cells, 15 cm dishes of 90% confluent cells in fed or under 72 h of serum depletion conditions were washed with 1x PBS. The cells were then lifted and transferred to a 2 ml tube, with an equivalent of 2.5% of the total cells reserved for further processing of the whole cell fraction. An insulin syringe (U-100 29G x ½”, Comfort point) was used to homogenize the cells with 5 strokes. The homogenate was centrifuged at 1000 g for 2 min at 4°C, after which the procedure continued as mentioned above. To confirm successful lysosomal protein precipitation, 2 µl and 10 µl of whole brain or cell samples and IP, respectively, were used for immunoblotting to detect LAMP1 and Cathepsin B (lysosomal markers), Golgin-97 (Golgi marker), Calreticulin (Endoplasmic reticulum marker), and VDAC (mitochondrial marker)

### Sample preparation for proteomic analysis of LysoIP samples

Eluates from LysoIPs and aliquots of the matched starting lysates were processed as described previously^13^. In brief, proteins were solubilized by addition of SDS to a final concentration of 2% (w/v), followed by sonication using the Bioruptor Plus (Diagenode) and heating for 10 min at 95°C. After reduction and alkylation, proteins were precipitated by cold-acetone precipitation. The resulting pellets were resuspended in digestion buffer (1 M Guanidinium Chloride in 100 mM HEPES pH 8.0) and digested by consecutive addition of LysC (3 h at 37°C) and trypsin (Promega, 16 h at 37°C). The obtained digested peptides were acidified and desalted using a Waters Oasis HLB μElution Plate 30 μm (Waters) according to the manufacturer’s instructions. The desalted peptides were dissolved in 5% (v/v) acetonitrile, 0.1% (v/v) formic acid to a peptide concentration of approximatively 1 μg/μl and spiked with iRT peptides (Biognosys) before analysis using LC-MS/MS. A detailed protocol can be found here^63^.

### Proximity labeling and sample preparation for BioID

Plasmids were generated using the pENTR Directional TOPO Cloning strategy (Thermo Fisher Scientific), enabling the integration of a PCR product into an entry vector, together with the Flp-In system (Invitrogen) for site-specific integration into the HEK-Flp cell line. All BirA plasmids are available at Addgene. To generate stable cell lines, Flp-HEK293 cells were transfected using XtremeGENE9 (Sigma), following the instructions of the manufacturer. In short, the plasmid containing target sequence was added to a ratio 1:9 with pOG44 plasmid (Invitrogen, Flp-Recombinase Expression Vector), in Opti-MEM media without antibiotics. After pre-incubation for 15 min at RT, the solution was added dropwise to the cells (200,000 in 6-well plate). On the next day the cells were transferred to a new plate containing selection antibiotics (final concentration Blasticidin 15 µg/ml and HygromycinB 100 µg/ml). The medium was replaced with fresh antibiotics twice a week, until colonies were seen growing on plates, taking two to three weeks. The colonies were then trypsinized to separate the cells and allow even growth. A more detailed protocol can be found here^64^.

For the BioID experiments, Flp-HEK293 cells were seeded at the density of approximately 1.6×10^4^ cells/cm^2^ and incubated for 24 h to allow cell attachment to the culture dish. The expression of BirA* fusion proteins was induced by addition of tetracycline (solved in ethanol) exposing the cells to its final concentration of 1 µg/µl in total for 4 days. 24 h prior to cell harvesting, 50 µM biotin was added to the culture media. Cells were washed 3 x times with PBS and harvested by trypsinization (0.05% trypsin incubated for 2 min at 37°C; Trypsin-EDTA (Gibco). For each sample, a pellet corresponding to 20×10^6^ cells was collected and snap-frozen in liquid nitrogen.

Cell pellets were thawed on ice and resuspended in lysis buffer (50 mM Tris pH 7.5, 150 mM NaCl, 1 mM EDTA, 1 mM EGTA, 1% (v/v) Triton X-100, 1 mg/ml aprotinin, 0.5 mg/ml leupeptin, 250 U turbonuclease, 0.1% (w/v) SDS), followed by 1 h incubation (15 rpm at 4°C) (as reported by^65^). Samples were sonicated in a Bioruptor Plus for 5 cycles (30 s on, 30 s off) at high setting and centrifuged at 20,817 *g*, for 30 min at 4°C. Streptavidin coated Sepharose beads were acetylated with 10 mM Sulfo-NHS-Acetate for 30 min at RT. The reaction was quenched with 1 M Tris pH 7.5 (1:10 v/v) and the beads washed three times with PBS (2000 *g*, 1 min at RT). Cleared lysates were transferred to new tubes and 50 µl of acetylated beads added (incubation for 3 h, 15 rpm at 4°C). After removing the supernatant (2000 *g* for 5 min at 4°C), the samples were transferred to a Pierce Spin Column (Snap Cap, Thermo Fisher Scientific). Beads were washed on the column with lysis buffer, followed by three washes with 50 mM ammonium bicarbonate (AmBic, pH adjusted to 8.3). The sample was transferred to a 2 ml tube and 1 µg of LysC added and incubated at 37°C for 16 h at 500 rpm. The samples were centrifuged at 2000 *g* for 5 min at RT and the content transferred to Pierce Spin column. Following elution, 0.5 μg of trypsin was added to the AmBic elutions and digested for 3 h, 500 rpm and 37°C. Digested peptides were acidified with 10% (v/v) trifluoroacetic and desalted using Waters Oasis HLB μElution Plate 30 μm (Waters) according to the manufacturer’s instructions. The desalted peptides were dissolved in 5% (v/v) acetonitrile, 0.1% (v/v) formic acid to a peptide concentration of approximatively 1 μg/μl and spiked with iRT peptides (Biognosys) before analysis using LC-MS/MS. A detailed protocol is described on protocols.io^66^.

### Proteomics data acquisition

Samples were reconstituted in MS Buffer (5% acetonitrile, 95% Milli-Q water, with 0.1% formic acid) and spiked with iRT peptides (Biognosys, Switzerland). Peptides were separated in trap/elute mode using the nanoAcquity MClass Ultra-High Performance Liquid Chromatography system (Waters, Waters Corporation, Milford, MA, USA) equipped with a trapping (nanoAcquity Symmetry C18, 5 μm, 180 μm x 20 mm) and an analytical column (nanoAcquity BEH C18, 1.7 μm, 75 μm x 250 mm)(BioID and LysoIP). 1 µl of the sample (∼1 μg on column) was loaded with a constant flow of solvent A at 5 μl/min onto the trapping column. Trapping time was 6 min. Peptides were eluted via the analytical column with a constant flow of 0.3 μl/min. Peptides were eluted using a non-linear gradient from 0% to 40% B in 90 min (for BioID) or 120 min (for all LysoIP, with exception of *CMV*-Cre). The total runtime was 115 min (BioID) or 145 min (for all LysoIP, with exception of *CMV*-Cre), including clean-up and column re-equilibration. The outlet of the analytical column was coupled directly to a Orbitrap Exploris 480 (Thermo Fisher Scientific, Bremen, Germany) system using the Proxeon nanospray source (BioID, and LysoIP, with exception of *CMV*-Cre). The peptides were introduced into the mass spectrometer via a Pico-Tip Emitter 360 μm outer diameter x 20 μm inner diameter, 10 μm tip (New Objective) heated at 300°C, and a spray voltage of 2.2 kV was applied. The capillary temperature was set at 300°C. The radio frequency ion funnel was set to 30%. MS acquisition parameters were set as follows: full scan MS spectra with a mass range of 350–1,650 m/z were acquired in profile mode in the Orbitrap with resolution of 120,000 FWHM. The filling time was set at a maximum of 60 ms with an AGC target of 3×10^6^ ions (for all LysoIP, with exception of *CMV*-Cre) or 5×10^5^ ions (BioID). DIA scans were acquired with 50 mass window segments of differing widths across the MS1 mass range. The default charge state was set to 3+. Higher energy collisional dissociation (HCD) fragmentation (stepped normalized collision energy; 25, 27, 30%) was applied and MS/MS spectra were acquired with a resolution of 30,000 FWHM with a fixed first mass of 200 m/z after accumulation of 3×10^6^ ions or after a filling time of 35 ms (for all LysoIP, with exception of *CMV*-Cre) or 40 ms (BioID)(whichever occurred first). Data were acquired in profile mode. For data acquisition and processing of the raw data Xcalibur 4.3 (Thermo) and Tune version 4.0 were used.

For whole brain LysoIPs (mockIP and *CMV*-Cre positive), peptides were separated using the nanoAcquity UPLC system (Waters) fitted with a trapping (nanoAcquity Symmetry C18, 5 µm, 180 µm x 20 mm) and an analytical column (nanoAcquity BEH C18, 1.7 µm, 75 µm x 250 mm). The outlet of the analytical column was coupled directly to a Thermo Q-Exactive HF using the Proxeon nanospray source. Solvent A was water with 0.1% formic acid, and solvent B was 80% (v/v) acetonitrile with 0.08% formic acid. Peptides were eluted via a non-linear gradient from 1% to 62.5% B in 131 min. Total runtime was 150 min, including clean-up and column re-equilibration. The same instrument settings as for the LysoIP (see above) were used, with the following exceptions: The RF ion funnel was set to 60%. The filling time was set at a maximum of 60 ms with an AGC target of 3×10^6^ ions. For data acquisition and processing Tune version 2.9 and Xcalibur 4.1 were employed.

### Parallel reaction monitoring (PRM)

Eleven peptides for the quantification of SLC45A1 were selected based on the most confident and consistently identified peptides of the LysoIP experiments of different mouse lines and the LysoIP of SH-SY5Y (Table S6). These peptides and their isotopically labeled version (heavy arginine (U-^13^C_6_;U-^15^N_4_) or lysine (U-^13^C_6_; U-^15^N_2_) at the C-term was added) were synthesized by JPT Peptide Technologies GmbH (Berlin, Germany) as SpikeTides L quality grade. Lyophilized peptides were delivered dried 10 nmol per peptide on a 96 well-plate and reconstituted in 20% (v/v) acetonitrile, 0.1% (v/v) formic acid with a final concentration of 0.05 nmol/µl and further pooled together in a ratio 1:1. The peptides mixture was used to make aliquots of 1 pmol/µl in 5% (v/v) acetonitrile in 50 mM AmBic and stored at -20°C. An aliquot of the pooled peptides, corresponding to approximatively 150 fmol per peptide, was analyzed by both DDA and DIA LC-MS/MS and used for assay generation using Spectrodive v.11.10.2 (Biognosys AG, Schlieren, Switzerland). 0.2 µg/µl LysoIP samples (Stock 1 µg/µl) were spiked with iRT (1:10, 0.5 µl) and JPT peptide mix (pooled stock 1:100, 1 µl) to a volume of 10 µl MS buffer (5% acetonitrile, 95% Milli-Q water, with 0.1% formic acid).

Peptides (5 µl) were separated using a nanoAcquity UPLC M-Class system (Waters, Milfors, MA, USA) with trapping (nanoAcquity Symmetry C18, 5 µm, 180 µm x 20 mm) and an analytical column (nanoE MZ HSS C18 T3 1.8 µm, 75 µm x 250 mm). The outlet of the analytical column was coupled directly to an Orbitrap Fusion Lumos (Thermo Fisher Scientific, Waltham, MA, USA) using the Proxeon nanospray source. Solvent A was water with 0.1% (v/v) formic acid, and solvent B was acetonitrile with 0.1% (v/v) formic acid. Peptides were eluted via the analytical column with a constant flow of 0.3 μl/min. During the elution step, the percentage of solvent B increased nonlinearly from 0% to 40% in 30 min. Total runtime was 60 min, including clean-up and column re-equilibration. PRM acquisition was performed in a scheduled fashion for the duration of the entire gradient (after instrument calibration in an unscheduled mode) using the “tMSn” mode with the following settings: resolution 120,000 FWHM, AGC target 3×10^6^, maximum injection time (IT) 250 ms, isolation window 0.4 m/z. For each cycle, a “full MS” scan was acquired with the following settings: resolution 120,000 FWHM, AGC target 3×10^6^, maximum injection time (IT) 10 ms, scan range 350 to 1650 m/z. For data acquisition and processing of the raw data Xcalibur 4.3 (Thermo) and Tune version 4.0 were used. Peak group identification was performed using Spectrodive and manually reviewed. Quantification was performed using a spike-in approach using the ratio between endogenous (light) and reference (heavy) peptides.

### Proteomics data processing for DIA data

Raw files were processed using Spectronaut Professional v15 to v18 (Biognosys; LysoIP mouse v18; LysoIP SH-SY5Y v17; BioID v15). Raw files were searched by directDIA search with Pulsar (Biognosys) against the mouse UniProt database (Mus musculus, release 2016_01 for LysoIP mouse; Homo sapiens, release 2016_01 for BioID and LysoIP SH-SY5Y) with a list of common contaminants appended, using the default settings. For quantification, the default BGS factory settings were used, except for the following settings: proteotypicity filter = only protein group specific; major group quantity = median peptide quantity; major group top N = OFF; minor group quantity = median precursor quantity; minor group top N = OFF; data filtering = *Q*-value percentile with fraction = 0.2 and imputing strategy = global imputing; normalization strategy = global normalization; row selection = identified in all runs (complete); modifications (only for BioID) = biotin_K. Differential abundance testing was performed in Spectronaut using a paired t-test between replicates. *P*-values were corrected for multiple testing multiple testing correction with the method described by Storey^67^. Here, also a Tmem119-Cre was included as another microglia cre, and included in the deposited data but not used for further analysis. The candidates and protein report tables were exported from Spectronaut and were used for further analysis using R v.3.6 and RStudio server 2024.04.0. Candidates and protein reports tables are available on PRIDE with the identifier MSV000096018 (LysoIP mouse), MSV000096053 (BioID), MSV000096063 (LysoIP SH-SY5Y).

### Logistic regression classifier for detecting LysoIP enriched proteins

To identify LysoIP enriched proteins, we trained a logistic regression binary classifier using an adapted version of the approach described in^65^. The classifier was trained using known core lysosomal proteins (Table S1B) as positive class, and nuclear chromatin associated proteins (defined by nuclear proteins from uniprot, Table S1C) as the negative set. Nuclear chromatin associated proteins are not expected to localize to lysosomes under homeostatic conditions. To distinguish between these two classes, we performed prediction using an enrichment score derived from multiplying the mean log2 ratio and the negative logarithm of the Q-value obtained from Spectronaut differential protein abundance analysis performed against control IPs. Before analysis, any missing data points were removed from the dataset. To assess the performance of the binary classifier, and optimize its parameters, a 10-fold cross-validation approach was adopted. The dataset was randomly partitioned into ten subsets, with nine subsets used for training and one subset for validation in each iteration. This process was repeated thirty times, and results were averaged using the mean value to ensure the robustness of the results. Logistic regression was employed as the classification method using the caret package in R^68^. To determine an optimal threshold for classification, the false positive rate (FPR) was set at 0.05. The threshold yielding an FPR closest to the target value of 0.05 was selected as the final classification threshold. Following model training and threshold selection, the classifier was applied to predict the class labels of additional proteins not used in the training process. The enrichment score and class labels for the new data were provided as input to the trained model. The pROC package was employed for ROC analysis^69^.

### Differential protein abundance between LysoIPs from different Cre lines

To test for differential protein abundance between LysoIPs performed from different Cre-lines, we extracted protein quantities from the Spectronaut report table for the 777 LysoIP-enriched proteins. Protein quantities were log2 transformed and the median quantity of core lysosomal proteins (Table S1B) were subtracted from each sample to account for differences in cell number and LysoIP efficiency. Differential protein abundance was determined using an ANOVA test applied to the normalized protein quantities using limma^70^. Correction for multiple testing was performed using the Benjamini Hochberg method. In order to estimate effect sizes, we calculated the difference between the mean log2 protein quantity in a given Cre line and the mean across all the Cre lines. Proteins that had an adjusted *P*-value < 0.05 and a maximum absolute log2 difference from the mean of all Cre lines > 1 were considered as differentially abundant.

### Comparison of protein abundance in LysoIPs and mRNA expression

To compare cell type specific abundance of LysoIP-enriched proteins and their corresponding mRNA expression in scRNASeq data^27^, the following approach was implemented. For each protein/gene pair, we computed the rank of expression across different cell types. The rank describes cell types that have the highest (high rank) and lowest (low rank) expression or abundance for each specific protein or gene. For LysoIP proteome data, a relative measure for protein abundance was used, described as the difference between the protein abundance in each cell type and the mean of the abundance of that specific protein across all cell types considered. For scRNA-Seq the normalized transcript abundance was quantified per cell cluster as the mean transcript levels across individual cells within each cluster, reflecting gene expression profiles of specific cell populations in the mouse brain. After computing a rank for each gene, to assess the similarity between protein and mRNA abundance for each gene, we calculated the rank-based Euclidean distance and derived a similarity score. First, for each gene/protein the Euclidean distance for each gene was computed as the square root of the sum of squared differences between the two ranks. To assign cell-type specificity for each gene/protein, the cell type with the highest rank was identified, to assign specificity. We then computed similarity between mRNA and protein levels by using the formula:

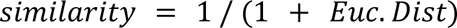

To assess the significance of the observed similarity between protein and mRNA abundance, we performed a randomization test. For each of 100 iterations, the ranks of mRNA abundance were randomly shuffled for all genes, while maintaining the ranks of protein abundance. In each iteration, the similarity measure was computed for each gene/protein using the same approach as described above. This process was repeated 100 times to generate a distribution of similarity scores under random conditions, which was later compared to the observed similarity score for each gene.

### Metabolite and lipid extraction and analysis

0.5×10^6^ of SH-SY5Y cells were seeded for 72 h in complete DMEM medium containing 10% heat-inactivated fetal bovine serum (Hi-FBS). For serum starvation, the cells were washed twice with 1x DPBS, then cultured in DMEM with 1% glutamine and 1% penicillin/streptomycin for 72 h.

Metabolites were extracted as described in^13^ and as described on protocols.io^71,72^. Briefly, metabolites were extracted in 80% methanol with isotopically labeled amino acids and stored at -80°C until analysis by LC-MS/MS. Metabolomics analysis was conducted as described on protocols.io^73^. Polar metabolites were profiled by LC/MS using a pHILIC column and the ID-X tribrid mass spectrometer. For data-dependent MS/MS (MS2) acquisition, pooled samples were used. Unbiased differential analysis was performed using Compound Discoverer 3.3 (CD), utilizing both local and online databases for MS1 and MS2-dependent annotation of metabolites and MS1-dependent quantitation of metabolites. Isotopically labeled amino acids were used for internal standardization. Following unfiltered, untargeted detection of features by CD (Table S10-Sheet 1), all features were manually filtered for the following criteria: 1) at least 1 annotation match with mzVault, mzCloud or local mass list library, 2) retention time of 3 min or greater and 3) correct CD peak integration. Following filtering, metabolites were manually annotated with putative identities as well as potential isomeric identities, if applicable, based on mzVault, mzCloud and local mass list library (Table S10-Sheet 2). Ultimately, post-filtering and post-manual annotation metabolites were used for generation of the untargeted volcano plot. Targeted quantitation of metabolites of interest was performed using TraceFinder 5.1 (TF). Retention times and m/z for metabolites of interest were collected from CD, and TF peak integration was performed, followed by manual correction of peak integrations when necessary. Ultimately, post-integration and post-correction peak areas were used for generation of the targeted metabolite quantitation plots.

Lipids were extracted as described in^13^ and as described on protocols.io^74^. Briefly, lipids were harvested with 2:1 (v/v) chloroform:methanol for 1 h, and further extracted after the addition of a 0.9% saline solution for 10 min. The chloroform phase was collected, dried in speed vacuum at RT, and stored at -80°C. Samples were reconstituted in a 13:6:1 acetonitrile:isopropanol:water (v/v/v) solution before analysis by LC/MS/MS. Lipidomics analysis was conducted as described on protocols.io^75^. Lipids were profiled by LC/MS using a C18 column and the ID-X tribrid mass spectrometer. Unbiased lipid annotation was performed using LipidSearch 4.2 (LS), utilizing MS1 and MS2-dependent annotation of lipids. Unbiased differential analysis was performed using Compound Discoverer 3.3 (CD), utilizing retention times, chemical formulas, and annotations from LS for MS1-dependent quantitation of lipids. Avanti Splash Lipidomix with labeled lipids was used for internal standardization. Following unfiltered, untargeted detection of features by CD (Table S7-Sheet 1), quantified features were defined as being significantly enriched or depleted by fold change greater than 2 and *P*-value less than 0.05 when comparing KO and rescued cells. These features were then manually filtered for the following criteria: 1) correct CD peak integration and 2) CD retention time match to LS. Following filtering, lipids were manually annotated with putative identities based on LS annotation, dependent on MS1, MS2 and retention time (Table S7-Sheet 2). For LS-annotated PGs, -H and +NH_4_ MS2s were manually inspected for definitive distinction between isomeric BMPs and PGs. BMPs were indicated by the dominant presence of monoacylglycerols in +NH_4_ MS2, and PGs were indicated by the dominant presence of diacylglycerol in +NH_4_ MS2. In cases where +NH_4_ MS2 was not acquired for definitive distinction of isomeric BMP and PG, the annotation BMP/PG was used. Ultimately, unfiltered lipids (Table S7-Sheet 1) were used for generation of the untargeted volcano plot, and manual annotations (Table S7-Sheet 2) were used for labeling of BMP and PG in the volcano plot. Targeted quantitation of lipids of interest was performed using TraceFinder 5.1 (TF). Retention times and m/z for lipids of interest were collected from LS and CD, and TF peak integration was performed, followed by manual correction of peak integrations when necessary. Ultimately, post-integration and post-correction peak areas were used for generation of the targeted lipid quantitation plots.

### Mouse cortical neuron dissection

Mouse cortical neurons were isolated from E16-17 mouse embryos following a protocol adapted from^76^. 10x Dissociation Media (DM) was prepared using Hanks’ Balanced Salt Solution (HBSS) without Mg^++^ and Ca^++^ (Gibco) including 100 mM MgCl_2_ (Sigma), 100 mM HEPES (Sigma), 10 mM kynurenic acid (Sigma). The solution was adjusted to a pH of 7.2 and stored at -20°C. Prior to use, 50 ml of 10x DM was diluted in 450 ml HBSS to obtain 1x DM. Following euthanasia, the amniotic sacs were removed from the pregnant mouse and placed on a sterile, ice-cold dissection dish. The embryos were decapitated, and brains were dissected under a microscope. The cortices were isolated and transferred to a cold 1x DM solution. Meninges were carefully removed, and cortices were collected for dissociation. Cortices were digested in a papain/DNAse mixture (Sigma) for 10 min at 37°C. The tissue was then washed with Neurobasal complete (NBc) media, and dissociated using a P1000 pipette, and filtered through a 0.4 µm cell strainer to remove undissociated tissue. NBc was prepared using a plain Neurobasal medium (Thermo Fisher Scientific) supplemented with B27 (Thermo Fisher Scientific), GlutaMax (Thermo Fisher Scientific), and penicillin/streptomycin (Thermo Fisher Scientific). The media was stored at 4°C and used for neuronal cultures. Plates were coated with Poly-D-lysine (Thermo Fisher Scientific) (5 mg/ml) and mouse laminin (Thermo Fisher Scientific) (0.5–2.0 mg/ml) the night before dissection. After coating, plates were incubated overnight at 37°C. The following day, plates were washed with sterile water and plain Neurobasal media before cells were plated. Dissociated neurons were plated at a density of 0.5×10^6^ cells per well in 12-well plates and incubated at 37°C with 5% CO₂. Cells were allowed to adhere for 4 h before media replacement.

### Immunofluorescence assays in cells and tissues

A total of 20,000 to 50,000 cells were plated on fibronectin-coated glass coverslips and fixed with 4% paraformaldehyde. The cells were then permeabilized with either 0.1x Saponin (Sigma) (for LAMP1 staining) or 0.1x Triton X-100 for 10 min at room temperature (RT) and blocked with 5% horse serum in PBS for 1 h at RT. This was followed by an overnight incubation at 4°C with primary antibodies diluted in 1% horse serum. Secondary antibodies (Alexa Fluor) were applied in 1% horse serum for 1 to 2 h. To stain the nucleus, 1:1000 Hoechst 33342 was used. Images were acquired using the Airyscan2 LSM980 microscope. For cryo-sectioning, immediately after euthanasia, brains were removed and fixed in 4% paraformaldehyde (VWR) overnight at 4°C. The brains were then transferred to 4% paraformaldehyde with 5% sucrose (Sigma) and left overnight at 4°C. Subsequently, the brains were placed in 15 ml of 30% sucrose in a 15 ml falcon tube to allow sinking at 4°C. The brains were then embedded in O.C.T (optimal cutting temperature) compound (Tissue-Tek), frozen, and stored at -80°C. For immunofluorescence staining, 10 to 20 µm sections were permeabilized with 0.3% Triton X-100 (Sigma) for 30 min. Sections were blocked in 5% horse serum and 0.3% Triton X-100 for 2 h. Primary antibodies: rat anti-LAMP1, mouse anti-HA, rabbit anti-HA, rabbit anti-NeuN, mouse anti-GFAP, rabbit anti-MBP, and rabbit anti-TMEM119 (details provided in the key resources table) were applied in 1% horse serum (Thermo Fisher Scientific) and 0.3% Triton X-100 for 24 h. Secondary antibodies were then applied in 1% horse serum and 0.3% Triton X-100 for 2 h. The sections were covered with coverslips using Vectashield (Vector Laboratories). Images were acquired using the Airyscan2 LSM980 with ZEN imaging software.

U2OS cells were grown on autoclaved coverslips (Carl Roth, YX03.1), coated with Poly-D-Lysine. Coverslips were placed individually in 12-well plates (Lab solute, 7696791), and 50×10^3^ cells were seeded per well. Transient transfection was performed by pre-mixing 1 µg DNA and 3 µg PEI (polyethylenimine, MW 25 kDa) in 100 µl OptiMEM (without serum and antibiotics). For a detailed protocol on how to prepare the transfection, follow protocols.io^77^. The transfection mix was incubated for 15 min at RT and added to the wells dropwise, either 20 µl (for PLEKHB1) or 30 µl per well (all other protein candidates). Incubation time was optimized to 6 h (CBLN3, TMEM256), 12 h (HOMER2, CBLN3, TMEM256, SLC45A1, ENPP5) or 24 h (HOMER2, EDIL3, PDGFA, PLEKHB1, CBLN3) for optimal transfection and replaced transfection media to DMEM and incubated further for 24 h (CBLN3, TMEM88B, TMEM256) or 48 h (for all protein candidates). Prior fixation, the cells were incubated with 75 nM Lysotracker (Red DND-99) for 1 h at 37°C. Cells were washed three times with PBS, fixed in 4% formaldehyde (v/v) in PBS for 10 min and incubated 5 min with DAPI (4’,6-Diamidino-2-Phenylindole, Dihydrochloride, 0.02 μg/μl in PBS) at RT, then washed 3 times with PBS. Coverslips were mounted in Permafluor mounting medium using glass slides (041300, Menzel) and dried at RT overnight. All samples were stored at 4°C in the dark until further analysis by microscopy. Images were acquired with a ZEISS Axio Imager (Z2 using a Plan-Apochromat 100x/1.4o Oil M27 Objective) with ZEN v.3.2 imaging software. Instrument settings were as follow: DAPI filter excitation wavelength 353 nm and emission 465 nm; Green fluorescence protein (GFP) filter excitation wavelength 488 nm and emission 509 nm; Lysotracker (DsRed) filter excitation wavelength 578 nm and emission 589 nm. The light source was a HXP 120V with intensity set to 25.9% for DAPI, 80% for GFP and 80% for LT, with the exposure time set to 23 ms for DAPI, 600 ms for GFP and 150 ms for LT. Co-localization was determined using Fiji with the plugin BIOP JACoP (parameter threshold: Otsu, on cropped ROIs^56,57^; to calculate the Manders coefficients^78^.

### Protein analyses and immunoblotting

Lysates were separated using SDS–PAGE (Novex wedgewell Tris-Glycine Mini Gels, Thermo Fisher Scientific) at 120 V. The proteins were then transferred to PVDF (Sigma) membranes at 40 V for 2 h. After the transfer, membranes were blocked with 5% BSA in TBST (Tris-buffered saline with Tween-20) for 1 h. They were then incubated overnight at 4°C with primary antibodies diluted in 5% bovine serum albumin (BSA) (Sigma) in TBST. Primary antibodies are listed in the key resources table. Following this incubation, membranes were washed three times with TBST for 5 min each. Next, they were incubated with secondary antibodies, diluted 1:3000 in 5% Non-Fat Dry milk, for 1 h at RT. After a final three washes with TBST, the membranes were developed using the ECL2 western blotting substrate (Thermo Fisher Scientific) to visualize the proteins.

Lysed BioID samples were sonicated in a Bioruptor Plus (4 cycles 60 s on, 30 s off) and boiled 5 min at 95°C in 1x Laemmli sample buffer (Bio-Rad), containing 5% □-Mercaptoethanol (v/v). Samples were then separated on an SDS-PAGE (precast 4–20% gel, Bio-Rad) at 100 V for 1.5 h and transferred in a TransBlot Turbo chamber (Bio-Rad) for 30 min to a nitrocellulose membrane (transfer buffer containing methanol and SDS). The membrane was then blocked for 1 h at RT with 3% BSA in TBST, and incubated 60 min at RT with Step-Tactin HRP (horse radish peroxidase) 1:100 dilution. Afterwards the membrane was washed three times with TBST for 5 min each. Proteins were detected using Pierce ECL Western Blotting Substrate for 1 min before being imaged on a molecular imager (ChemiDoc XRS+, Bio-Rad) and analyzed by Image Lab.

### Electron microscopy

For mouse study, immediately following euthanasia, the brains were harvested. The cortices were fixed overnight at 4°C in a solution of 4% paraformaldehyde (PFA) and 2% glutaraldehyde (provided by the Stanford University Cell Sciences Imaging Core Facility, RRID:SCR_017787). The next day, the cortices were transferred to a solution containing 4% PFA, 2% glutaraldehyde, and 5% sucrose, and left overnight at 4°C. Afterward, the tissues were placed in 15 ml of 30% sucrose solution in Falcon tubes, at 4°C to allow sinking. Once the tissues had sunk, they were embedded in O.C.T compound (TissueTek), frozen, and stored at -80°C. Coronal sections, 200 µm thick, were cut using a Leica VT-1000 vibratome (Leica). For cell-based studies, 0.2×10^6^ SH-SY5Y wild-type, *SLC45A1*-KO, and rescue cells were seeded onto Aclar coverslips (provided by Stanford University Cell Sciences Imaging Core Facility, RRID:SCR_017787). The cells were washed with 1x DPBS and treated with serum-depleted DMEM. To distinguish lysosomes from other vacuoles, the cells were treated with 0.2 g/100 ml BSA-20-nm gold conjugate as described by Rabinowitz et al.^79^. Lysosomes were counted as long as they looked similar to BSA containing vesicles. The samples were then transferred to the Stanford University Cell Sciences Imaging Core Facility (RRID:SCR_017787) for further processing and electron microscopy imaging.

### RNA extraction and qPCR from cells and tissues

Total RNA was extracted using miRNeasy Mini kit according to the manufacturer’s protocol (Qiagen). For cells, 1×10^6^ cells were lysed by adding 700 ml QIAzol lysis reagent. For tissues, 25 mg was used for RNA isolation. cDNA was synthesized with 0.5-1 μg of total RNA using High-Capacity cDNA Reverse Transcription Kit (Applied Biosystems, or RevertAid First Strand cDNA Synthesis Kit Thermo Scientific, according to the manufacturer’s instructions). qPCR was performed in a Bio-Rad CFX96 Real-Time System using Syber green (FastStart Universal SYBR Green Master) following manufacturer’s instructions. All samples were assessed to the levels of β-Actin expression as an internal control. qPCR data were assessed and reported according to the 2^-ΔΔCt^ method.

### Flow cytometry for LAMP1 staining

Cells were lifted using trypsin and then centrifuged. The cell pellet was resuspended in 1 ml of 3% formaldehyde in PBS and incubated for 10 min at 37°C. The cells were then centrifuged and resuspended in 90% ice-cold methanol while vortexing at low speed to prevent clump formation. The cells were incubated on ice for 30 min, briefly vortexing every 10 min. After centrifugation (∼1000 g, 3 min) and decanting the supernatant, the cells were washed with PBS. Following another centrifugation, the cells were resuspended in 1-2 ml of 3% BSA in PBS and incubated for 30 min at 37°C. After centrifugation, the cells were resuspended in 3% BSA in PBS containing a 1:1000 dilution of LAMP1 antibody (Rabbit-anti human LAMP1, Cell Signaling Technology,). The mixture was incubated for 1-2 h at 37°C, with intermittent vortexing. After centrifugation, the cells were resuspended in the wash buffer (PBS + 1% BSA + 0.05% Tween-20) and centrifuged again. The cells were then resuspended in 3% BSA in PBS containing a 1:1000 dilution of Alexa Fluor-647 and incubated for 30 min to 1 h at 37°C. The cells were washed twice with wash buffer, centrifuged, and finally resuspended in wash buffer. The intensity of LAMP1 was measured using flow cytometry. A detailed protocol is described here^80^.

### Reactive Oxygen Species (ROS) measurement

250×10^3^ cells were seeded in a 12-well plate with complete medium. After 72 h, the cells were washed twice with 1x DPBS, followed by the addition of DMEM without FBS for 72 h. Subsequently, the cells were treated with 5 µM CellROX-Green (Invitrogen) for 30 min. The cells were then washed with PBS, trypsinized, and the intensity of CellROX was measured by flow cytometry.

### Cathepsin B activity assay

20×10^3^ cells were seeded in 96 wells-plate. After 72 h, cells were washed twice with 1xDPBS and DMEM medium without FBS were added for another 72 h. Cells were then washed with 1xDPBS and assay was performed following manufacturer’s instructions. Briefly, Reconstituted Magic Red Substrate (MR-RR2) (Immunochemistry technology) with 50 μl DMSO was then diluted 1:10 with diH_2_O. Magic Red staining solution was added at 1/25 (20 μl staining solution to 480 μl cells forming a final volume of 500 μl) to cells for 30 min incubation at 37°C. Cells were immediately imaged by fluorescent microscopy (Keyence, BZ-X710).

### Co-immunoprecipitation (co-IP)

15×10^6^ SH-SY5Y cells expressing Flag-GFP-SLC45A1 or wildtype cells were seeded in 15 cm dishes. After 72 h, the cells were washed with 1x PBS and then lysed with 1 ml of 1:10 diluted extraction buffer (1x extraction buffer: 50 mM HEPES at pH 7.4, 40 mM NaCl, and 2 mM EDTA, 1% Triton X-100, 1.5 mM sodium orthovanadate (NaVO_4_), 30 mM sodium fluoride (NaF), 10 mM sodium pyrophosphate (Na_4_P_2_O_7_), 10 mM sodium β-glycerophosphate) with protease inhibitor cocktail for 1 h on a shaker at 4°C. The lysates were centrifuged for 10 min at maximum speed. 100 µl of the supernatant was kept as input and added 20 µl 5x loading dye. The remaining 900 µl was added into a new tube containing 25 µl of pre-washed (1:10 diluted extraction buffer) Anti-Flag magnetic beads (Thermo Fisher Scientific). The lysates were incubated overnight at 4°C with rotation. The beads were washed seven times with 1 ml of 1:10 diluted extraction buffer, changing the tubes every two washes. After the last wash, 50 µl 1x loading dye was added to beads. Proteins were separated using SDS–PAGE (Thermo Fisher Scientific) following protein analyses and immunoblotting. The list of antibodies used is provided in the key resources table.

### Measurement of lysosomal acidity

To measure the acidity of lysosomes in wildtype, *SLC45A1*-KO and rescue cells expressing SLC45A1, we used Fluorescence indicator reporting pH in Lysosomes (FIRE-pHLy)^54^. Cells were transduced with pFUGW-FIRE-pHLy vector, which was a gift from Aimee Kao (Addgene plasmid #170774). After selection with puromycin, 0.5×10^6^ cells were seeded with complete media. After 3 days, cells were washed twice with 1xDPBS and incubated with serum depleted DMEM for 72 h for live cells imaging. Images were acquired using the Airyscan2 LSM980 microscope. Images were processed and intensity was measured using ImageJ (Fiji) software.

### Seahorse assay

To assess changes in mitochondrial function and cellular metabolism, cells underwent the Agilent Seahorse XFe Cell Mito Stress test (Agilent, Santa Clara, CA, USA), which measures mitochondrial function through oxygen consumption rate (OCR). The test was performed according to the manufacturer’s instructions. 80,000 cells were seeded in Seahorse XFe96 microplates and incubated at 37°C with 5% CO_2_ for 48 h. For the iron supplementation experiment, cells were seeded and treated with 0.2 mg/ml ferric ammonium citrate (FAC) (Fisher Scientific) for 48 h to assess the effects of increased iron levels. For the ROS inhibition experiment, cells were seeded and treated with 250 nM Liproxstatin-1 (Lip1) (Cayman Chemical) for 48 h to evaluate the impact of reactive oxygen species (ROS) inhibition on cell viability and stress responses. On the day of the assay, the growth medium in the cell culture microplate was replaced with a pre-warmed (37°C) assay medium including Seahorse XF base medium (Agilent Technologies), 1 mM Pyruvate (Thermo Fisher Scientific), and 20 mM glucose. The microplate was incubated in a non-CO_2_ incubator at 37°C for 1 h to allow the medium to reach temperature and pH equilibrium. Modulating agents were prepared in assay medium and injected as final concentration: 1 µM oligomycin, 50 µM 2,4-Dinitrophenol (DNP) as an uncoupler, and 0.5 µM rotenone. OCR was measured and analyzed using the Seahorse XFe Report Generator. At the end of the assay, cells were lysed, and total protein was extracted and measured for normalization.

### Sucrose and invertase treatment

A total of 15×10^6^ SH-S5Y5 cells were seeded in 15 cm cell culture dishes and incubated for 3 days under fed condition. Then cells were subjected to serum starvation conditions for 72 h. After 48 h of serum deprivation, sucrose was added to the culture medium at a final concentration of 50 mM for an additional 24 h, making the total incubation time 72 h. Following the sucrose (Sigma) treatment, the cells were washed thoroughly with 1x DPBS to remove residual sucrose. Following the washing step, cells were incubated with 0.5 mg/ml of invertase from *Saccharomyces cerevisiae* (Sigma) in serum-free DMEM for 1 h. This enzyme catalyzed the hydrolysis of accumulated sucrose into glucose and fructose, facilitating the enrichment of hexoses within lysosomes. After the invertase treatment, lysosomal isolation was performed using the LysoIP to capture intact lysosomes. The isolated lysosomal fractions were then subjected to benzoyl-hexose derivatization, allowing for precise quantification and analysis of hexose content within the lysosomes.

### Benzoyl-Hexose derivatization

This protocol was adopted from^46^. To derivatize hexoses to benzoyl-hexose, 200 µl of a 1:1 ethyl acetate:water mixture was added to the purified lysosomes. Samples were vortexed for 5 min and then centrifuged at 16,000 g for 5 min. After removing the upper layer and transferring the bottom layer to a new tube, 20 µl of 1 M potassium phosphate dibasic (K_2_HPO_4_) (Sigma), 20 µl of 8 M NaOH (Fisher Scientific), and 10 µl of benzoyl chloride (Sigma) were added. The mixture was vortexed for 5 min and then briefly spun down. Next, 10 µl of 1.4 M phosphoric acid (H_3_PO_4_) and 500 µl of ethyl acetate (Fisher Chemical) were added, followed by vortexing for 3 min and centrifugation at 16,000 g for 5 min. Subsequently, 400 µl of the ethyl acetate upper layer was transferred to a new tube, and 100 µl of 1 M NaOH was added. The mixture was vortexed for 2 min and centrifuged for 5 min at 16,000 g. Then, 100 µl of the ethyl acetate upper layer was transferred to a new tube. This step was repeated with the addition of 100 µl of 1 M NaOH to the original tube, followed by 2 min of vortexing, 5 min of centrifugation at 16,000 g, and the transfer of 100 µl of the ethyl acetate upper layer to the same tube, for a total of 200 µl. The samples were then dried in a speed vacuum and reconstituted in 50 µl of a 9:1 acetonitrile:water mixture containing 100 mM ammonium formate (NH_4_HCO_2_) (Sigma). Samples were vortexed for 10 min and centrifuged at maximum speed for 10 min. For analysis, we monitored for parent 718 m/z [M^+^NH_4_]^+^ and product ions 231, 105, and 579.1.

### Data acquisition and analysis of derivatized hexose/glucose

Derivatized hexose/glucose was separated on an Eclipse Plus C18 column (4.6 x 100 mm, 3.5 µm) (Agilent) and connected to a 1290 II LC system. An Ultivo triple quadruple (QQQ) mass analyzer equipped with an LC-ESI probe was coupled to the LC system. An external mass calibration was performed using the standard calibration mixture every seven days. An injection volume of 3 µl was used for each sample with fast polarity switching. Mobile phase A was composed of 10 mM ammonium formate and 0.1% formic acid in LC/MS grade 60:40 water:acetonitrile. Mobile phase B was composed of 10 mM ammonium formate and 0.1% formic acid in 90:10 isopropanol:acetonitrile. The chromatographic gradient was the following: linear increase from 50%-100% B for 3 min; isocratic elution from 3-6 min with 100% B; fast linear decrease from 100%-50% B from 6-6.1 min; 50% B hold for 1.9 min. The flow rate was set to 0.400 ml/min, and the column compressor and autosampler were held at 30°C and 4°C, respectively.

The mass spectrometer parameters were as follows: the spray voltage was set to 3.6 kV in positive mode and 2.5 kV in negative mode, and the gas temperature and the sheath gas flow were held at 250°C and 400°C, respectively. The gas and sheath gas flows were 11 l/min and 12 l/min, respectively. The nebulizer was maintained at 28 psi.

The precursor-product ion pairs (m/z) used for MRM of the compounds were the following:

| Compound | Precursor ion | Product ion | Fragmentor Voltage | Collision Energy |
| --- | --- | --- | --- | --- |
| Benzoyl-Hexose/Glucose | 718.0 | 231.0 | 100 | 11 |
| Benzoyl-Hexose/Glucose | 718.0 | 105.0 | 100 | 37 |
| Benzoyl-Hexose/Glucose | 718.0 | 579.1 | 100 | 5 |

High-throughput annotation and relative quantification of hexose/glucose were performed using qualitative analysis software for MassHunter acquisition data, along with QQQ quantitative analysis (Quant-My-Way) software. Hexose/glucose intensities were validated by manually checking peak alignment in the Qualitative software to ensure retention times and MS/MS spectra matched the characteristic fragmentation of standard compounds. For further confirmation of accurate identification and quantification, two transitions for each compound were analyzed to verify consistent relative responses. Quantification of all hexose/glucose compounds was done using the MRM method and retention times, with raw peak areas exported to Microsoft Excel for further analysis.

### Data preparation and statistics

All quantitative graphs were generated in GraphPad Prism 9. Two-tailed independent t-tests, ordinary one-ANOVA, or Two-way ANOVA were used for statistical comparisons in Prism 9. If other tests were used, they were indicated in the legend. Figure diagrams and schematics were generated using BioRender and used with a permissive license.

### Data and materials availability

All MS data, source code and raw data will be archived in general-purpose repositories with free access following their peer review and upon publication.

